# The influence of axonal beading and undulation on axonal diameter mapping

**DOI:** 10.1101/2023.04.19.537494

**Authors:** Hong-Hsi Lee, Qiyuan Tian, Maxina Sheft, Ricardo Coronado-Leija, Gabriel Ramos-Llorden, Ali Abdollahzadeh, Els Fieremans, Dmitry S. Novikov, Susie Y. Huang

## Abstract

We consider the effect of non-cylindrical axonal shape on axonal diameter mapping with diffusion MRI. Practical sensitivity to axon diameter is attained at strong diffusion weightings *b*, where the deviation from the 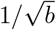 scaling yields the finite transverse diffusivity, which is then translated into axon diameter. While axons are usually modeled as perfectly straight, impermeable cylinders, the local variations in diameter (caliber variation or beading) and direction (undulation) have been observed in microscopy data of human axons. Here we quantify the influence of cellular-level features such as caliber variation and undulation on axon diameter estimation. For that, we simulate the diffusion MRI signal in realistic axons segmented from 3-dimensional electron microscopy of a human brain sample. We then create artificial fibers with the same features and tune the amplitude of their caliber variations and undulations. Numerical simulations of diffusion in fibers with such tunable features show that caliber variations and undulations result in under- and over-estimation of axon diameters, correspondingly; this bias can be as large as 100%. Given that increased axonal beading and undulations have been observed in pathological tissues, such as traumatic brain injury and ischemia, the interpretation of axon diameter alterations in pathology may be significantly confounded.

## 1. Introduction

Diffusion MRI (dMRI) probes tissue microstructure at the mesoscopic scale and enables estimation of cellular-level features, such as axon diameter and cellular size (Novikov et al., 2019). Identifying the alteration of axon diameter in the brain white matter helps to characterise the tissue pathology for early diagnosis and monitoring treatment responses in, for example, multiple sclerosis (Lovas et al., 2000; DeLuca et al., 2004; Huang et al., 2016, 2020) and amyotrophic lateral sclerosis (Cluskey and Ramsden, 2001; Tandan and Bradley, 1985).

This signal sensitivity to the axon diameter, further weighted by the axon volume, gives a strong preference to the tail of axon diameter distribution (Burcaw et al., 2015; Veraart et al., 2020), providing the interpretation of axonal diameter mapping (ADM) results in the brain using dMRI with strong gradients (Assaf et al., 2008; Alexander et al., 2010; Huang et al., 2016, 2020; Fan et al., 2020; Veraart et al., 2020).

So far, ADM has been validated in perfectly straight cylinders and relatively straight, thick axons (Drobnjak et al., 2016; Nilsson et al., 2017; Andersson et al., 2022). In white matter, ADM is confounded by a number of known factors: the non-Gaussian (time-dependent) diffusion in extra-axonal space, relevant at low-to-moderate diffusion weightings, both transverse (Burcaw et al., 2015; Fieremans et al., 2016; Lee et al., 2018) and along axons (Novikov et al., 2014; Fieremans et al., 2016; Lee et al., 2020b); and orientational dispersion (Veraart et al., 2020; Fan et al., 2020), observed even in highly aligned fiber regions such as corpus callosum (Ronen et al., 2014; Lee et al., 2019) and spinal cord (Grussu et al., 2016; Jespersen et al., 2018). Besides, inter-compartmental water exchange between the intra- and extra-axonal space is assumed to be slow and negligible at clinical diffusion times (Deoni et al., 2008; Lampinen et al., 2017).

To reduce the confound from the extra-axonal space, one can apply strong enough diffusion weighting to suppress the extra-axonal signal (Veraart et al., 2020). To factor out the fiber orientation dispersion, diffusion signals are directionally averaged — known as powder-averaging (Jespersen et al., 2013) or the spherical mean technique (SMT) (Kaden et al., 2016) — for each diffusion weighting *b*, and the deviation from the 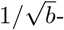scaling at high *b* yields an estimate of axon diameter (Veraart et al., 2020; Fan et al., 2020).

In this work, we consider in detail the previously unexplored confounds, stemming from axons being not ideal straight cylinders — namely, *caliber variations* or *beading* (Budde and Frank, 2010; Lee et al., 2020a), and *undulations* (changes in local axon direction) (Nilsson et al., 2012; Brabec et al., 2020; Lee et al., 2020a). The impact of realistic geometries such as caliber variation and undulation on ADM has been first noticed in mouse brain axons extracted from 3-dimensional electron microscopy (EM) (Lee et al., 2020a), where ADM is only accurate for straight axons at shorter time scales *δ <* 10 ms. However, the effect of caliber variation and undulation on ADM has not been separated and quantified in human brain using realistic tissue substrates with tunable cell features.

Recently, a 1.4 petabyte EM volume of human temporal lobe tissue and the adjacent subcortical white matter was made publicly available (Shapson-Coe et al., 2021). This dataset can serve as a valuable resource for building numerical phantoms for the characterization and validation of ADM in the human brain white matter.

Here, we segment 76 myelinated axons (33.5-189.3 *μ*m in length, Figure 1) in subcortical white matter of this human brain EM sample using U-Nets (Tian et al., 2020) and generate undulating, beaded fibers of circular cross sections with tuned caliber variations and undulations (Figure 3a) similar to those observed in real axons on EM. We then calculate intra-axonal diffusion signals using Monte Carlo (MC) simulations of diffusion in segmented axons and artificial fibers. For fibers of the same scaling of caliber variations and undulations, we generate intra-axonal diffusion signals by taking their volume-weighted sum. Diffusion in the extra-axonal space and cerebrospinal fluid (CSF) is approximated by anisotropic or isotropic Gaussian diffusion, but is generally inessential as we perform parameter estimation at high *b*. To reveal the effect of axonal features on axonal diameter mapping using spherical mean signals, we test ADM models in two scenarios:

**Figure 1:**
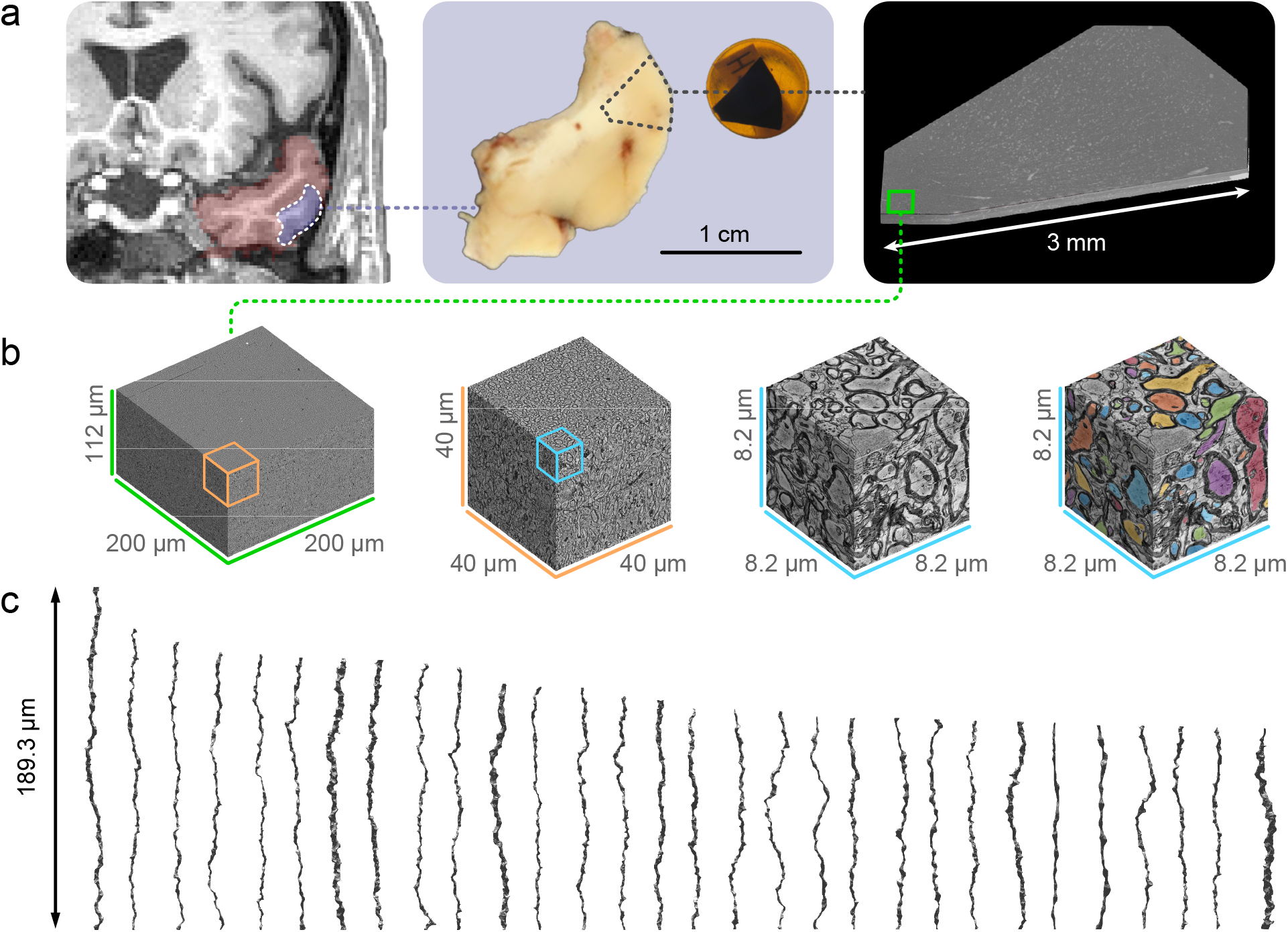
(a) Non-lesional brain tissue (purple) was obtained from the left middle temporal gyrus (red) of a 45-year-old female undergoing surgery for hippocampal sclerosis, drop-fixed in glutaraldehyde/paraformaldehyde fixative, stained with osmium tetroxide, and embedded in resin (Shapson-Coe et al., 2021). The tissue was subsequently scanned with a high-resolution multi-beam scanning electron microscope. (b) The 200×200×112 *μ*m^3^ subset of the EM volume corresponding to subcortical white matter (green rectangle in the right panel of (a)) was segmented using 2d and 3d U-Nets. (c) The segmented intra-axonal space of myelinated axons longer than 33.5 *μ*m were aligned along the z-axis, resulting in 76 axons ranging from 33.5 to 189.3 *μ*m long. The figure (a) is adapted from (Shapson-Coe et al., 2021) with permission from bioRxiv.

(i) First, we fit Callaghan et al.’s (1979) model (isotropically distributed impermeable cylinders) to the spherical mean of intra-axonal signals. We call this model a single-compartment SMT (Section 2.6.1, Figure 4a). The effect of axonal features on axonal diameter mapping is evaluated in isolation, excluding the extra-axonal space and CSF.
(ii) Further, we add a CSF component and an extra-axonal space locally aligned with axons in the model, where Gaussian diffusion is assumed (Assaf et al., 2008). We refer to this model as a multi-compartmental AxCaliber-SMT (Section 2.6.2, Figure 4b) (Fan et al., 2020). This model is fit to spherical mean of the combined intra-axonal, extra-axonal and CSF signals for the full range of *b*-values. The time-dependent non-Gaussian effects in the extra-axonal space in this approach can only be neglected when operating at sufficiently high *b* values. By fully modeling the diffusion in the intra- and extra-axonal space, and CSF, we are able to evaluate the effect of axonal features on not just axon diameter but also other tissue parameters, such as intra-axonal and CSF volume fractions, and extra-axonal diffusivity.

MC simulations in both cases show that caliber variations and undulations result in under- and over-estimation of the axon diameter, respectively.

## 2. Methods

All the data are publicly available in the literature (Shapson-Coe et al., 2021) following the ethical standards of Harvard University. This article does not contain any studies with human participants and animals performed by any of the authors.

### 2.1 Human brain EM segmentation

Non-lesional brain tissue was obtained from the anterior portion of the left middle temporal gyrus of a 45-year-old female during surgery for resection of an epileptogenic focus in the left hippocampus (Shapson-Coe et al., 2021) (Figure 1a). The pathological evaluation showed hippocampal sclerosis, whereas the brain tissue sampled from the anterior portion of the middle temporal gyrus showed no diagnostic abnormality recognized. The tissue sample was trimmed to 2×3×0.2 mm^3^ and scanned with a multi-beam serial-section scanning electron microscope (Sigma, Carl Zeiss) in 4×4 nm^2^ pixels and 33 nm slice thickness. The 200×200×112 *μ*m^3^ subset of subcortical white matter was down-sampled to 32×32 nm^2^ pixels and 33 nm slice thickness and segmented using 2d and 3d U-Nets (Tian et al., 2020) (Figure 1b), which was initially trained on ground truth segmentation of myelin and intra-axonal space in EM images of the mouse corpus callosum (Lee et al., 2019). The voxel size was further down-sampled to 64×64×64 nm^3^ after segmentation. The segmented intra-axonal spaces of myelinated axons longer than 33.5 *μ*m were aligned along the z-axis, resulting in 76 axons ranging from 33.5-189.3 *μ*m long (Figure 1c).

### 2.2 Axon morphology analysis

The inner axonal radius along individual axons was estimated by using the equivalent circle radius *r*, defined as the radius of an equivalent circle with the same cross-sectional area as the intra-axonal space (West et al., 2016; Lee et al., 2019; Abdollahzadeh et al., 2019; Lee et al., 2020a). The caliber variation of individual axons was defined as the coefficient of variation of the radius (Lee et al., 2019, 2020a)

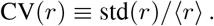

The axon radius estimated by MR is heavily weighted by thick axons (Neuman, 1974; Burcaw et al., 2015). Diffusion weighting, introduced by applying magnetic field gradients, leads to spatial variations of the Larmor frequency across an axon cross-section. For practically relevant case of wide-pulsed gradients, when gradient duration *δ* exceeds the time to diffuse across axon radius *r*, signal attenuation can be interpreted as transverse relaxation (Lee et al., 2018), − ln 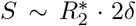, occurring during the net pulse duration 2*δ*, with the rate 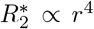 that is strongly sensitive to the radius (Neuman, 1974; van Gelderen et al., 1994). This signal sensitivity is further weighted by the axon volume ∝ *r*^2^, leading to the effective MR axon radius *r*_eff_ calculated based on the equivalent circle radius *r* (Burcaw et al., 2015; Sepehrband et al., 2016; Lee et al., 2020a; Veraart et al., 2020):

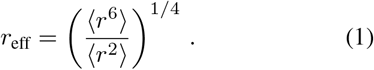

This is the histological reference to be compared with ADM results.

To quantify axonal undulations, the axonal skeleton was built by connecting the center of mass of each intraaxonal cross section along individual axons. The undulation amplitude *w*_0_ and wavelength *λ* were calculated using a simplified single harmonic model for the axonal skeleton (Lee et al., 2020a, 2021).

### 2.3 Undulating, beaded fibers similar to actual axons

To mimic axonal geometries based on their most relevant features and understand the impact of these features on axonal diameter mapping, we created artificial fibers of circular cross sections with the same undulations and caliber variations as in actual axons (Lee et al., 2020b). We retrieved the axonal skeletons of axons from EM and replaced each cross section with a circle of the same cross-sectional area of the actual axons, i.e., in the equivalent circle radius *r*. Furthermore, tuneable axonal geometries were created by taking fibers with varying undulation amplitude *p*_*w*_ · *w*_0_ and coefficients of variation of the radius *p*_*r*_ · CV(*r*), each scaled from *p*_*w*_ = 0% (no undulation) to 100% (strong undulation) and *p*_*r*_ = 0% (no caliber variation) to 100% (strong caliber variation). The undulation wavelength *λ* and axon volume of each axon were kept unchanged.

### 2.4 Monte Carlo (MC) simulations

MC simulations were implemented in CUDA C++ for diffusion in the 3-dimensional micro-geometry of intraaxonal space from the selected 76 axons and their fiber derivatives aligned in the z-direction. The cell shapes were described by voxelized geometries based on the voxel size (64 nm)^3^ of EM segmentation. 100,000 random walkers per fiber were employed, diffusing for 1.1×10^5^ steps with a duration of 2.8×10^−4^ ms and a step length of 58 nm for each step (Fieremans and Lee, 2018; Lee et al., 2020a,b, 2021). The intrinsic diffusivity was set to *D*_0_ = 2 *μ*m^2^/ms (Dhital et al., 2019; Novikov et al., 2018; Veraart et al., 2019). To prevent discontinuities within the axonal geometries, the top and bottom faces of each fiber were extended by reflective copies using mirroring boundary conditions. The diffusion signal resulting from a pulsed-gradient spin-echo sequence was calculated, with pulse duration *δ* = 10 ms, inter-pulse interval Δ = 20 ms, eight b-values *b* = [1, 2, 3, 5, 7, 12, 17, 26] ms/*μ*m^2^, and 60 gradient directions per b-shell. The MC simulations took 10 days on an NVIDIA V100 GPU core. The normalized simulated signal of the *i*-th axon in gradient direction *ĝ* was denoted as 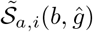 with the tilde indicating signal derived from the MC simulations. The normalized spherical mean signal 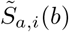 of the *i*-th axon was calculated by averaging diffusion weighted signals 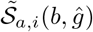 in 60 directions over each *b*-shell.

### 2.5. Signal generation

To demonstrate the effect of axonal shape on diameter mapping, we generated normalized spherical mean signals in one compartment (i.e., intra-axonal space) or multiple compartments (i.e., intra-axonal space, extra-axonal space, and CSF). In this study, all diffusion signals *S* were normalized, such that the non-diffusion-weighted signal *S*_0_ ≡ 1.

#### 2.5.1. Spherical mean signal in one compartment

The intra-axonal signal 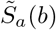 was created by randomly choosing 38 out of 76 axons’ fiber derivatives (using the same scaling factor combination of undulations and caliber variations, (*p*_*w*_, *p*_*r*_) in Section 2.3) over 500 iterations for bootstrapping and calculating the volume-weighted sum of their simulated spherical mean signals 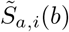:

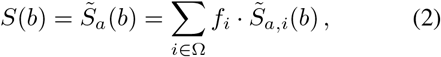

where Ω is a list of the chosen axons in each iteration, *f*_*i*_ is the volume fraction of the *i*-th fiber in the list Ω, such that Σ_*i*∈Ω_ *f*_*i*_ = 1.

#### 2.5.2. Spherical mean signal in multiple compartments

To create signals similar to those in white matter, we combined signals in two additional compartments (extra-axonal space and CSF) with the simulated spherical mean signal 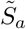 in intra-axonal space in Section 2.5.1:

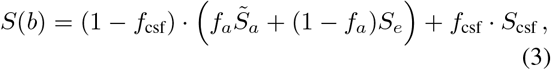

where *f*_*a*_ and *f*_csf_ are the relaxation-weighted volume fractions of the intra-axonal space and CSF. *S*_*e*_ and *S*_csf_ are the normalized spherical mean signals in the extra-axonal space and CSF, where diffusion is modeled as axi-symmetric Gaussian ellipsoids and free diffusion, respectively (Callaghan et al., 1979; Jespersen et al., 2013; Kaden et al., 2016; Fan et al., 2020):

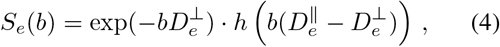

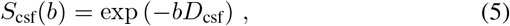

where

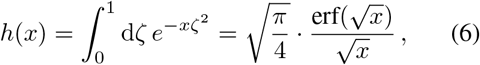

erf(·) is the error function, 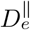 and 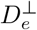 are the extraaxonal diffusivities along and transverse to the axonal segments, respectively, and the CSF diffusivity *D*_csf_ was fixed at 3 *μ*m^2^/ms. The simulated intra-axonal signal 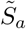 in each bootstrapping iteration was combined with the extra-axonal space and CSF signals, and the value of each parameter was randomly chosen in the range of *f*_*a*_ ∈ [0.5, 1], *f*_csf_ ∈ [0, 0.2], 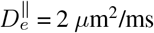, and 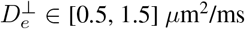.

We note that signal (3) with the extra-axonal contribution (4) assumes Gaussian extra-axonal diffusion within each fiber fascicle and neglects the time-dependent diffusion effects (Burcaw et al., 2015) that can further bias ADM at low *b*. Hence, its applicability is skewed towards large *b*, where *S*_*e*_ ≪ 1 is negligible.

### 2.6. Axonal diameter mapping

Axon diameter estimation using the spherical mean signals factors out fiber dispersion and simplifies the functional form in model fitting. Here we compared two approaches to performing axonal diameter mapping using the spherical mean signal (Figure 4). The first approach used the single-compartment SMT model for intra-axonal signals. The second approach used the multi-compartmental AxCaliber-SMT model for the combined signals of intra-axonal, extra-axonal and CSF signals.

#### 2.6.1. Single-compartment SMT, Figure 4a

Approximating real axons as a collection of cylindrical segments of length 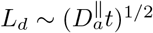 at diffusion time *t*, the normalized spherical mean signal of the intraaxonal space is given by (Callaghan et al., 1979; Jespersen et al., 2013; Kaden et al., 2016; Veraart et al., 2019; Veraart et al., 2020; Fan et al., 2020)

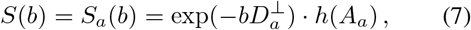

where *h*(*x*) is defined in Eq. (6),

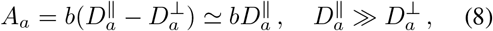

and 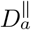 and 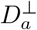 are the intra-axonal diffusivities along and transverse to the axonal segments, respectively. An axon radius of 1 *μ*m yields an estimate of 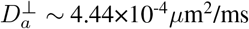 (Neuman, 1974) with *D*_0_ = 2 *μ*m^2^/ms at *δ* = 10 ms and Δ = 20 ms. In other words, 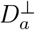 is smaller than 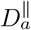 by three orders of magnitude, justifying the approximation in Eq. (8). This model is fit to the spherical mean signal of the intra-axonal space generated by the MC simulations (Eq. (2)) using nonlinear least squares with positive constraints for all parameters. This is a two-parameter fit with parameters 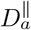 and 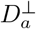.

The estimated 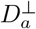 is translated into the MR-estimated axon radius in Neuman’s limit (Neuman, 1974; Veraart et al., 2020):

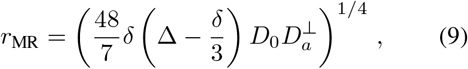

with intrinsic diffusivity *D*_0_ fixed at 2 *μ*m^2^/ms, matching the value in simulations. Eq. (9) is applicable in the wide pulse limit, i.e., *δ* ≫ *r*^2^/*D*_0_ (Neuman, 1974; van Gelderen et al., 1994). Given that the effective axon radius is about 1 *μ*m in histology (dominated by the tail of axon radius distribution, Eq. (1) and Figure 2a), the pulse width *δ* = 10 ms is indeed much longer than the correlation time *r*^2^/*D*_0_ ∼ 0.5 ms.

**Figure 2:**
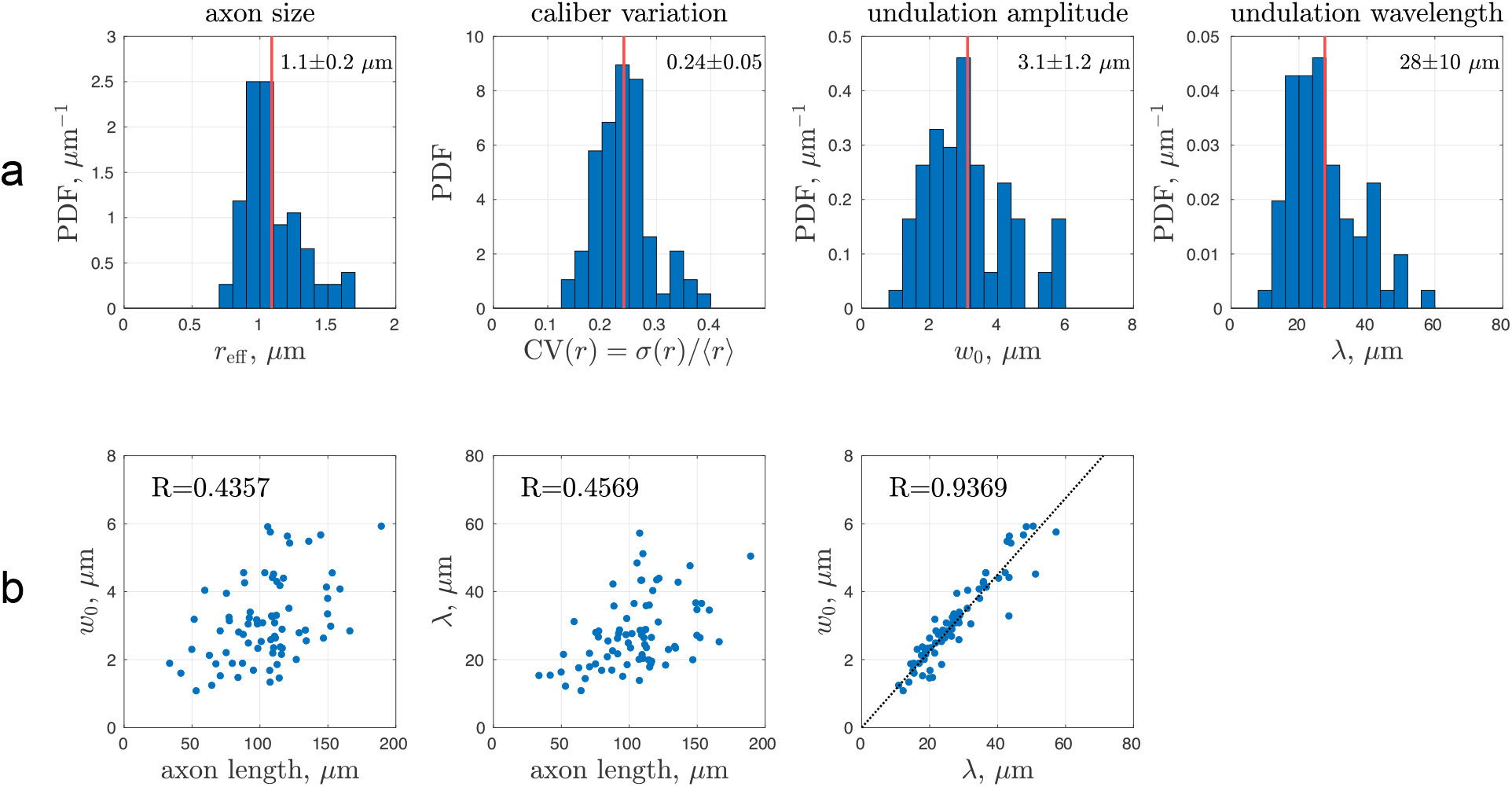
Features of myelinated axons in the human white matter sample. (a) Effective MR axon radius *r*_eff_ and caliber variation CV(*r*) were calculated based on the equivalent circle radius *r*, and undulation amplitude *w*_0_ and wavelength *λ* were calculated using a simplified single harmonic model for axonal skeletons. The mean (red line) and standard deviation were reported. (b) Undulation amplitude and wavelength were highly correlated, and both showed much lower correlations with axon length.

To evaluate the performance of axonal diameter mapping with and without noise, we fit the single compartment model in Eq. (7) to the noiseless signal 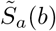 in Section 2.5.1 and its magnitude signal with Rician noise added, where the noise levels in the real and imaginary parts of the signal are both *σ* = *S*_0_/SNR (Gudbjartsson and Patz, 1995; Koay and Basser, 2006) with non-diffusion weighted signal *S*_0_ ≡ 1 and SNR = ∞ (no noise) or 100, respectively.

#### 2.6.2. Multi-compartmental AxCaliber-SMT, Figure 4b

To describe multiple compartments in white matter for both low and high *b*, we model the spherical mean signal as consisting of contributions from the intra-axonal space, extra-axonal space, and CSF (Kaden et al., 2016; Fan et al., 2020):

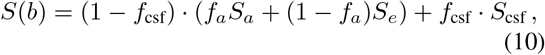

where *S*_*a*_, *S*_*e*_, and *S*_csf_ are the spherical mean signals described in Eqs. (7), (4), and (5), and time-dependent diffusion effects in the extra-axonal space are neglected. This is a six-parameter fit with parameters (*f*_*a*_, *f*_csf_, 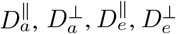), and *D*_csf_ fixed to 3 *μ*m^2^/ms. It is difficult to reliably fit six parameters due to the parameter degeneracy problem (Jelescu et al., 2016; Novikov et al., 2018), and thus 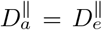 were both fixed at 1.2, 1.7, or 2 *μ*m^2^/ms to stabilize the fitting, leading to a four-parameter fit. The axon radius *r*_MR_ is again estimated based on 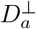 in Neuman’s limit in Eq. (9).

To evaluate the ADM performance, we fit the multi-compartmental model in Eq. (10) to the combined signal in Eq. (3) using nonlinear least squares with positive constraints for all parameters. The magnitude signals are composed of MC-simulated signals in intra-axonal space and added signals in the extra-axonal space and CSF with and without the Rician noise, where the noise levels in real and imaginary parts are both *σ* = *S*_0_/SNR with *S*_0_ ≡ 1 and SNR = ∞ (no noise) and 100, respectively.

### 2.7. Comparison of estimated axonal radii with histology

To evaluate the accuracy and precision of the model fitting, we compared the MR-estimated axon radius *r*_MR_ in Eq. (9) with the histological reference *r*_eff_ in Eq. (1). We calculated their offset and the normalized root mean square error (NRMSE), which is defined as the ratio of RMSE to the reference’s mean value. Similarly, for additional parameters in multi-compartmental AxCaliber-SMT model, we calculated the offset and NRMSE of the estimated parameters (*f*_*a*_, *f*_csf_, 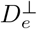) and the values used for signal generation, which served as the ground truth.

### 2.8. Data and code availability

The EM data can be downloaded on the Neuroglancer (Shapson-Coe et al., 2021). The simulation codes can be downloaded on the Github page.

## 3. Results

### 3.1. Single-compartment SMT

The spherical mean of the MC-simulated signals of diffusion within human brain axons segmented from EM (Figure 3b) yielded a radius estimate *r*_MR_ *<* 0.001 *μ*m, much smaller than the histological *r*_eff_ ≈1.1 *μ*m (Figure 2a). This bias may arise from axonal undulations or caliber variations. To determine the most relevant features contributing to this bias, we translated the real axons from EM into artificial fibers of circular cross sections with the same undulations and caliber variations, whose simulated signals were almost the same as those of real axons (Figure 3b and Figure S1), and then scaled these features to generate artificial fibers with varying caliber variation and undulation amplitude (Figure 3a).

**Figure 3:**
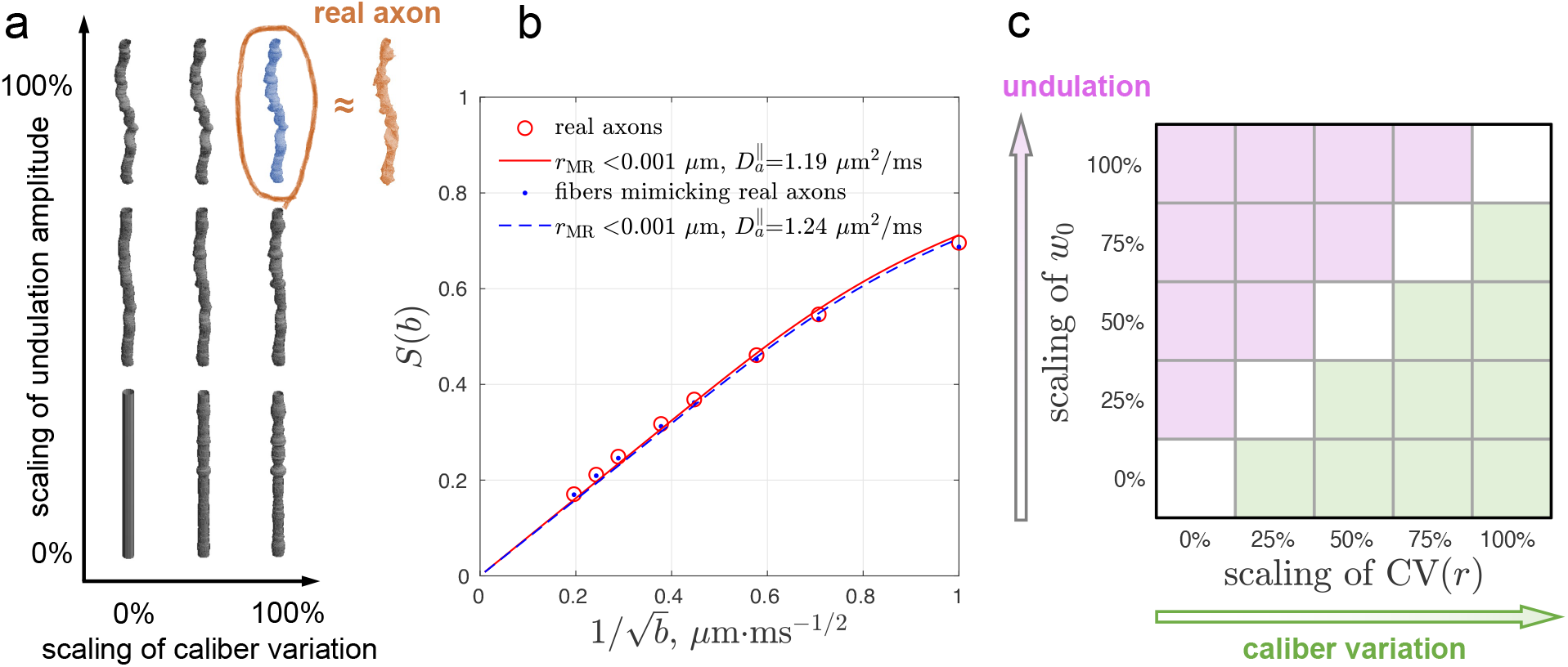
To mimic axonal geometries based on their most relevant features, we created (a) fibers of circular cross sections (blue) with the same undulations and caliber variations as real axons (red), and fibers of circular cross sections (gray) with undulation amplitudes *w*_0_ and coefficients of variation of radius CV(*r*) scaled from 0% to 100%. (b) The simulated spherical mean signals in real axons (Figure 1c) and axon-mimicking fibers (panel a, blue) were consistent, and their MR-estimated axon radii *<* 0.001 *μ*m were much smaller than the histological ground truth 1.1 *μ*m in Figure 2a. (c) In this study, fitting results were shown in a 5×5 grid. The upper left corner corresponded to results of fibers with strong undulations (magenta), and the lower right corner corresponded to results of fibers with strong caliber variations (green).

**Figure 4:**
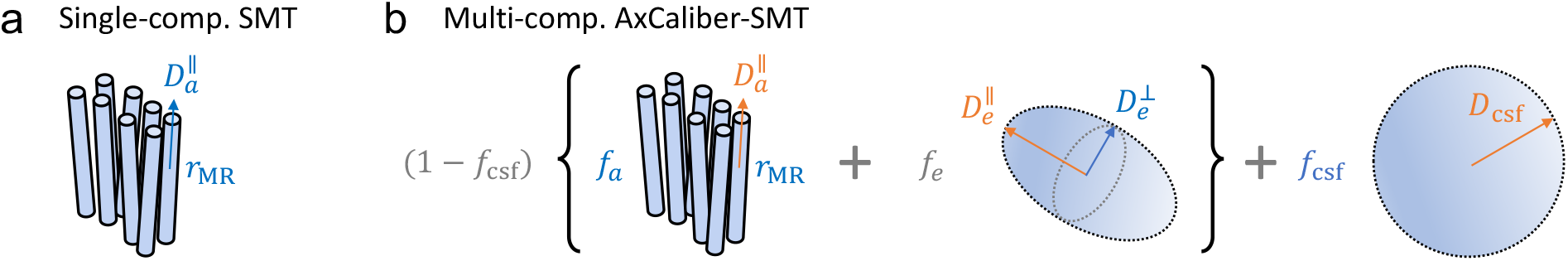
Biophysical modeling of the diffusion MRI signal for axonal diameter mapping. (a) Single-compartment SMT (Section 2.6.1) only includes the spherical mean of the intra-axonal signal in Eq. (7). We fit this model to the spherical mean MC-simulated signals of the intra-axonal space (Section 2.5.1). (b) Multi-compartmental AxCaliber-SMT (Section 2.6.2) includes the spherical mean of the intra-axonal space, extra-axonal space, and CSF signals in Eq. (10). We fit this model to the generated spherical mean signals of the intra-axonal space, extra-axonal space, and CSF (Section 2.5.2). In each model, estimated parameters are in blue, fixed parameters are in orange, and the dependent parameters are in gray, e.g., extra-axonal volume fraction *f*_*e*_ = 1 − *f*_*a*_.

For the case without noise, the simulated spherical mean signals in the artificially generated fibers led to radius estimates *r*_MR_ greater than the histological *r*_eff_ (overestimation) in the fibers with strong undulations, and radius estimates *r*_MR_ smaller than the histological *r*_eff_ (underestimation) in the fibers with strong caliber variations (Figure 5a). The estimated axial diffusivity 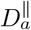 decreased with undulations and caliber variations.

**Figure 5:**
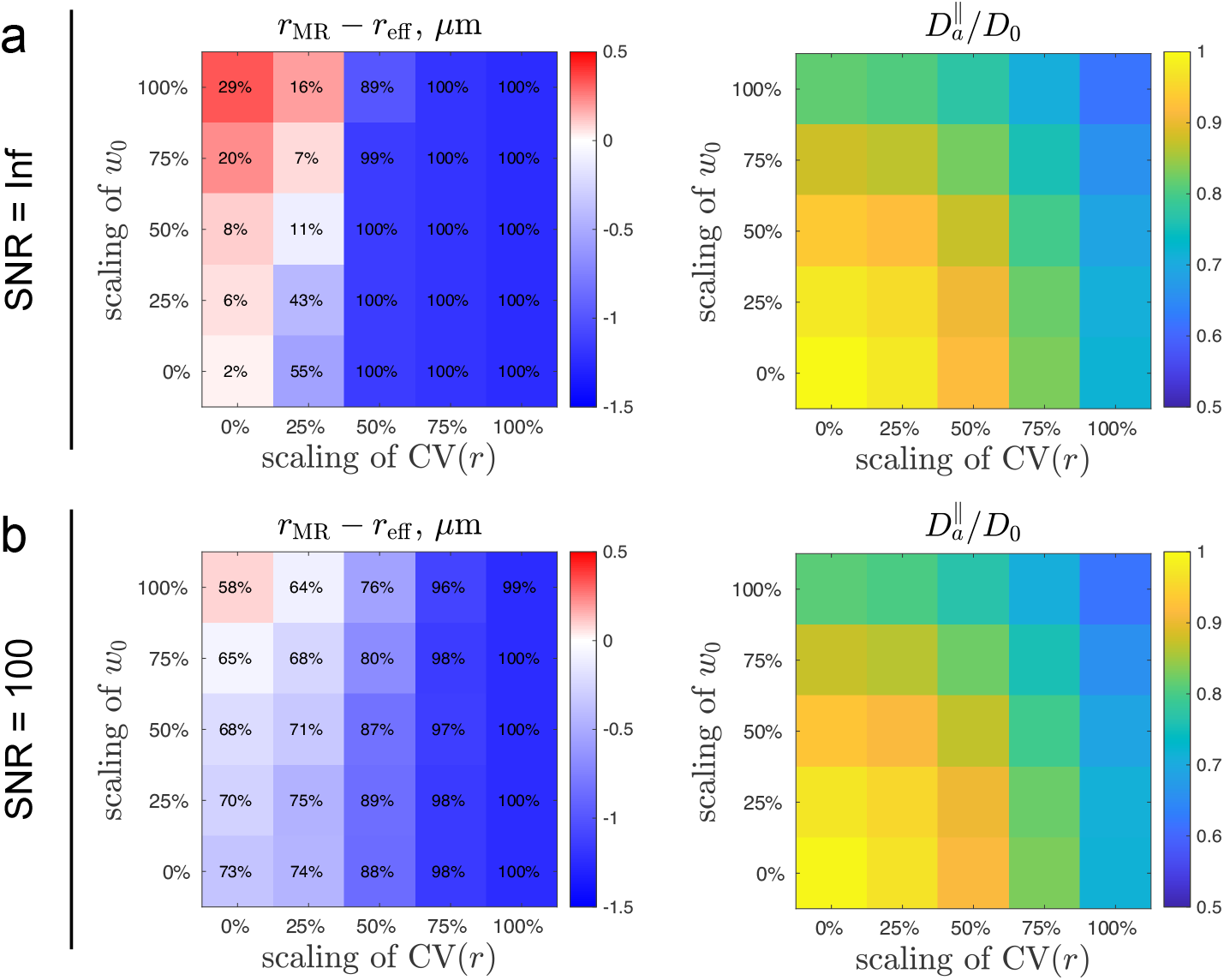
Axon size estimation based on diffusion simulations in undulating, beaded fibers reveals the effect of undulations and caliber variations on axonal diameter mapping, respectively. The single-compartment SMT model in Section 2.6.1 is fitted to intra-axonal signals in Section 2.5.1. The number in each pixel indicates the normalized root mean square error (NRMSE) between the estimation *r*_MR_ and histology *r*_eff_. When a parameter is underestimated, its NRMSE cannot exceed 100% due to the positive constraint in nonlinear least square fitting. (a) Simulations without adding the noise show that undulations lead to overestimation of axon size, and caliber variations lead to underestimation of axon size. Both undulations and caliber variations lead to a decrease in intra-axonal axial diffusivity. (b) Simulations with Rician noise (SNR = 100) show that Rician noise floor leads to underestimation of axon size in low precision, manifested by large NRMSE. The estimated intra-axonal axial diffusivity is relatively unaffected by the noise.

Simulations with added Rician noise (SNR = 100) showed that the axon diameter was generally underestimated due to the Rician noise floor, in both cases of undulations and caliber variations (Figure 5b). Furthermore, the precision was much lower due to the noise, which was manifested in the large values for NRMSE. In contrast, the estimated axial diffusivity decreased with undulations and caliber variations, and was relatively unaffected by the noise.

### 3.2. Multi-compartmental AxCaliber-SMT

To estimate axon size in white matter using diffusion data from all b-values, we combined the intraaxonal MC-simulated signals with axi-symmetric Gaussian signals representing the extra-axonal space and isotropic Gaussian signals representing the CSF. We fitted the multi-compartmental AxCaliber-SMT model in Eq. (10) to the combined signals in Eq. (3). The noiseless data led to overestimated axon size in axons with strong undulations and underestimated axon size in axons with strong caliber variations (Figures 6a, S2a, and S3a). In contrast, the combined signal with added Rician noise (SNR = 100) resulted in underestimated axon size due to the Rician noise floor, for both cases of undulations and caliber variations (Figures 6b, S2b, and S3b).

**Figure 6:**
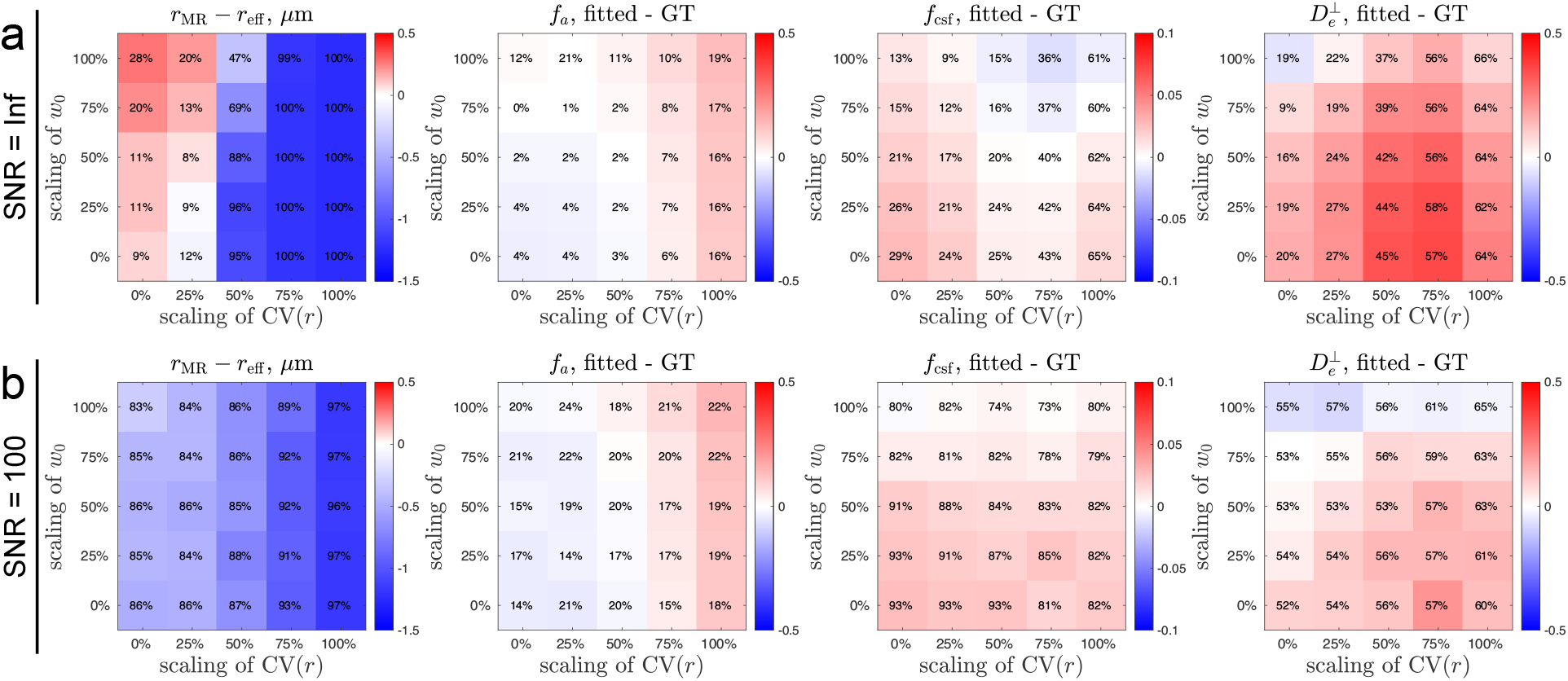
Axon size estimation based on diffusion simulations in undulating, beaded fibers reveals the effect of undulations and caliber variationson axonal diameter mapping, respectively. The multi-compartmental AxCaliber-SMT model 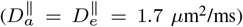 in Section 2.6.2 is fitted to the multi-compartmental signal in Section 2.5.2. The number in each pixel indicates the NRMSE between the estimation and ground truth. When a parameter is underestimated, its NRMSE cannot exceed 100% due to the positive constraint in nonlinear least square fitting. (a) Simulations without adding the noise show that undulations lead to overestimation of axon size, and caliber variations lead to underestimation of axon size. In contrast, the estimated intra-axonal volume fraction *f*_*a*_ shows the opposite trend; undulations lead to slight underestimation of *f*_*a*_, and caliber variations lead to overestimation of *f*_*a*_. In addition, both undulations and caliber variations lead to overestimation of CSF volume fraction *f*_csf_ and extra-axonal radial diffusivity 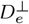. (b) Simulations with Rician noise (SNR = 100) show that the Rician noise floor leads to underestimation of axon size with low precision, manifested by large NRMSE. The bias in *f*_*a*_, *f*_csf_, and 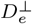 is similar to the noiseless result, and yet the precision is lower, i.e., larger NRMSE. The radial diffusivity 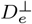of the extra-axonal space is in units of *μ*m^2^/ms. GT = ground truth.

The biases in estimated volume fractions (*f*_*a*_, *f*_csf_) and radial diffusivity in the extra-axonal space 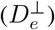 were relatively unaffected by the Rician noise, though with slightly higher values of NRMSE (Figures 6, S2, and S3). The intra-axonal volume fraction was slightly underestimated in axons with strong undulations and overestimated in axons with strong caliber variations. Estimated CSF volume fraction and extra-axonal radial diffusivity were both overestimated while fixing 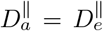 at 1.2 or 1.7 *μ*m^2^/ms, for both cases of undulations and caliber variations (Figures S2 and 6). In contrast, for 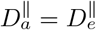 fixed at 2 *μ*m^2^/ms, the estimated CSF volume fraction was underestimated for both cases of undulations and caliber variations, whereas the extra-axonal radial diffusivity was under- and over-estimated due to undulations and caliber variations, respectively (Figure S3).

## 4. Discussion

In this work, we report the results of Monte Carlo simulations of diffusion inside axons segmented from EM images of human temporal subcortical white matter. Using the simulated diffusion signals, we study the performance of axonal diameter mapping using biophysical models of the diffusion MRI signal. To explore the effect of cellular-level features (i.e., undulations and beading) on MR-estimates of axon diameter, we create fibers of circular cross sections with varying scales of undulations and caliber variations based on those observed in the segmented axons and perform diffusion simulations in these fiber derivatives. Numerical simulations in 3-dimensional EM-based micro-geometries serve as a critical validation step for biophysical modeling of the diffusion MRI signal and provide important insights into the interpretation of quantitative biomarkers of axon size from diffusion MRI, particularly in fixed tissue, and their alterations in pathology.

### 4.1. Single-compartment SMT

Axonal diameter mapping in white matter may be affected by strong undulations and caliber variations in axonal shape, as is observed in the fixed human brain tissue imaged here by EM. Caliber variations have previously been considered to be a major contributor to the overestimation of axon size since the MR-measured effective radius in Eq. (1) is heavily weighted by the tail (large axons) of radius distribution (Burcaw et al., 2015; Sepehrband et al., 2016; Veraart et al., 2020); however, our simulations show that caliber variations lead to underestimation of axon size in the actual model fitting to the spherical mean signals. The variation of local axial diffusivity and axial kurtosis along the length of individual axons may play a role in explaining this result (Section 4.3).

For the case with added Rician noise (SNR = 100), the axon size is generally underestimated for axons with varying degrees of undulation and caliber variation. This can be understood based on an analysis of the resolution limit in axonal diameter mapping (Nilsson et al., 2017; Andersson et al., 2022). To evaluate the resolution limit, we first calculate the difference of the normalized spherical mean signals in Eq. (7) between infinitely thin cylinders (sticks) and those with finite ra-dius *r*:

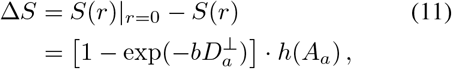

with *h*(·) and *A*_*a*_ defined in Eqs. (6) and (8). For thin fibers with radius ∼1 *μ*m, the intra-axonal radial diffusivity is 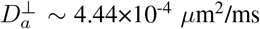 based on Neuman’s solution (9) at *δ* = 10 ms, Δ = 20 ms and *D*_0_ = 2 *μ*m^2^/ms. The 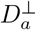 is much smaller than the intra-axonal axial diffusivity 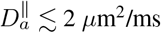, justifying the approx-imation 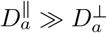. Given the above estimation of 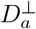, we approximate 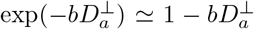 even at high b-values (e.g., up to 26 ms/*μ*m in this study), further simplifying the analytical form of signal difference:

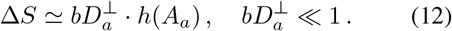

To observe significantly different diffusion signals between zero-radius fibers and finite radius ones, the z-score *z*(Δ*S*) of the signal difference is required to be larger than the z-threshold *z*_*α*_ for significance level *α* (Nilsson et al., 2017):

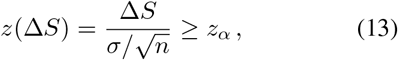

where the noise level *σ* is defined by the SNR ≡ *S*_0_/*σ* with *S*_0_ ≡ 1, and the number *n* of signal averages is the number of gradient directions per b-shell. Substituting Eqs. (9) and (12) into Eq. (13), we obtain the resolution limit for axonal diameter mapping using spherical mean signals (Nilsson et al., 2017):

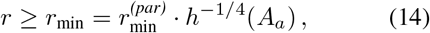

where 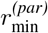 is the resolution limit for axonal diameter mapping by applying diffusion gradients transverse to highly aligned cylinders,

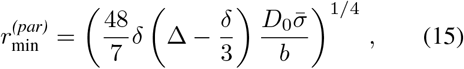

and 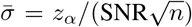. In the study, we have *D*_0_ = 2 *μ*m^2^/ms, 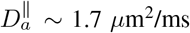, maximal b-value *b* = 26 ms/*μ*m^2^, *δ* = 10 ms, Δ = 20 ms, *n* = 60, and *z*_*α*_ = 1.64 for *α* = 0.05, yielding and estimate of resolution limit *r*_min_ ∼ 1.09 *μ*m at SNR = 100. The resolution limit *r*_min_ is smaller than the histological values *r*_eff_ ∼ 1.1 *μ*m in Figure 2a, explaining the high bias in the axon size estimation due to the Rician noise in Figure 5b. For very thick axons (e.g., *r >* 8 *μ*m for our sequence protocol), the values of 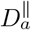 and 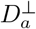 are comparable, and the resolution limit should be calculated by using the full functional form as in (Andersson et al., 2022). When diffusion signals are contributed by multiple compartments, the intra-axonal signal-to-noise ratio should be re-defined as SNR ≡ *f*_*a*_*S*_0_/*σ* with *f*_*a*_ the intra-axonal volume fraction. This leads to even larger 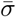 and subsequent increase of *r*_min_, making it even harder to estimate the size of thin axons.

In addition to the axon size, the estimated axial diffusivity 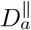 decreases in axons with strong undulations and/or caliber variations. This is expected based on previous simulation studies: Budde and Frank (2010) created fibers with periodic beads and showed that simulated axial diffusivity decreased with the strength of beading. Lee et al. (2020b) performed diffusion simulations in realistic axonal shapes from mouse brain EM and showed that axial diffusivity decreased mainly due to caliber variations. Lee et al. (2020a) performed diffusion simulations in undulating fibers of circular cross sections and showed the reduction of axial diffusivity due to undulations. In Section 4.2, we further demonstrate the correlation of estimated 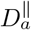 and these axon features.

### 4.2. Axial diffusivity is correlated with undulations and caliber variations

The measurement of axial diffusivity along axons is usually confounded by the fiber dispersion. Instead, estimating axial diffusivity 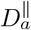 based on the spherical mean signal effectively factors out the effect of dispersion, revealing the actual value of axial diffusivity and its correlation with undulations and caliber variations.

In the following sections, we introduce the full functional form of axial diffusivity time-dependence due to undulations and caliber variations. However, the estimated axial diffusivity in axonal diameter mapping is an average along axonal segments (of length 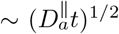) that may orient in slightly different directions due to undulations. The estimated axial diffusivity is not exactly the same as the axial diffusivity projected along the axon’s main axis. Further, the bias in axon diameter estimation can lead to bias in estimated axial diffusivity. Therefore, without the consideration of higher order effects (time-dependence), we only correlate the estimated axial diffusivity in axonal diameter mapping with the theoretical predictions due to undulations and caliber variations in the long time limit (*t* → ∞), respectively.

#### 4.2.1. The impact of undulations on axial diffusivity

To investigate the axial diffusivity time-dependence due to undulations, we evaluate the diffusivity along a simplified single harmonic, undulating fiber with no caliber variations. In the wide pulse limit of undulations, i.e., *δ* ≫ *t*_*u*_ ∼ *λ*^2^/*D*_0_, the axial diffusivity in a undulating thin fiber is given by (Appendix A)

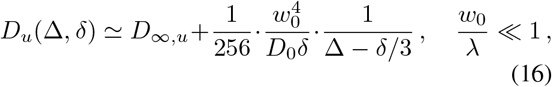

where

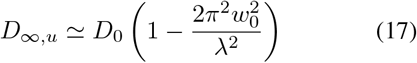

is the axial diffusivity at long time (Lee et al., 2020a). The approximation of small undulation is justified by the small value *w*_0_/*λ* ≈ 0.11 in real human axons (Figure 2). Interestingly, Eq. (16) indicated that the strength of diffusivity time-dependence along an undulating fiber is mainly affected by the undulation amplitude *w*_0_, but not the wavelength *λ*. Furthermore, in the wide pulse limit, the intra-axonal axial diffusivity due to undulations shares the same functional form of time-dependence in intra-axonal radial diffusivity (transverse to fibers) due to undulations (Lee et al., 2020a) and caliber variations (Neuman, 1974; Burcaw et al., 2015).

In simulations and diameter mapping of individual axons (Figure 5), the estimated axial diffusivity 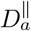 correlates well with the long-time prediction *D*_∞,*u*_ (17) due to undulations for axons with strong undulations (Figure 7a), though the correlation becomes weaker due to the Rician noise at SNR = 100 (Figure 7b).

**Figure 7:**
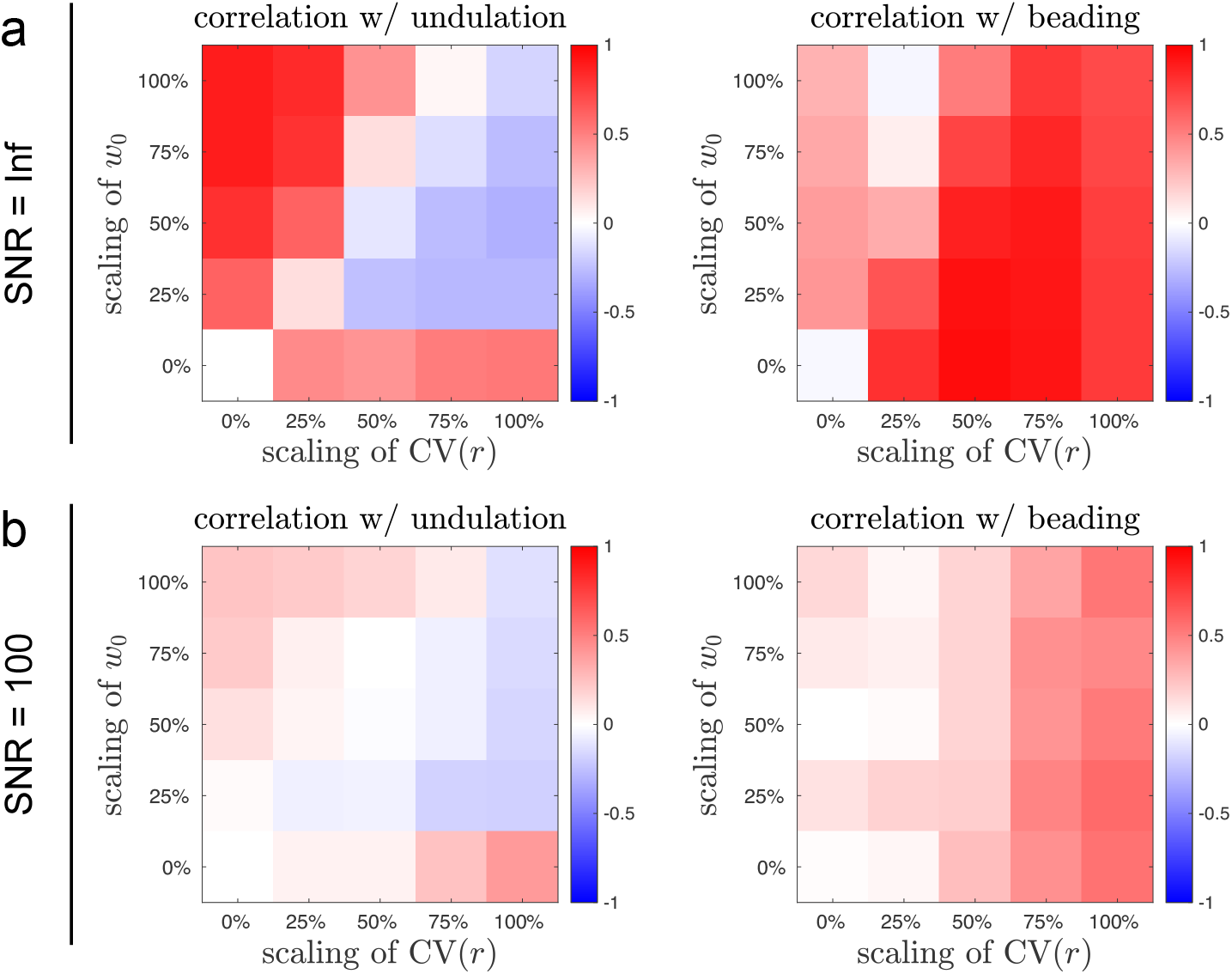
Intra-axonal axial diffusivity estimation based on diffusion simulations in undulating, beaded fibers reveals its correlation with undulations and caliber variations, respectively. The single-compartment SMT model in Section 2.6.1 is fitted to intra-axonal signals in Section 2.5.1. The color indicates the Pearson correlation coefficient between the estimated 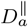 and the theoretical prediction due to undulations (*D*_∞,*u*_ in Eq. (17)) and caliber variations (*D*_∞,bead_ in Eq. (19)) in left and right panels, respectively. (a) Simulations without adding the noise show that the prediction *D*_∞,*u*_ due to undulations strongly correlates with undulations, and the prediction *D*_∞,bead_ due to caliber variations strongly correlates with caliber variations. (b) Simulations with Rician noise (SNR = 100) show similar trend in correlations with lower strength.

#### 4.2.2. The impact of caliber variations on axial diffusivity

Diffusion along neurites in the brain gray matter and white matter is restricted by caliber variations (beading), spines, shafts, branching, and other microstructural inhomogeneity (Hellwig et al., 1994; Glantz and Lewis, 2000; Shepherd et al., 2002; Morales et al., 2014; Lee et al., 2020b), whose long-range fluctuations are characterized by Poisson statistics and a structural exponent *p* = 0 in the power spectrum along neurites. At clinical diffusion times, diffusion in three-dimensional neurite structures effectively boils down to one-dimensional diffusion along neurites due to the coarse-graining over diffusion lengths longer than the typical length scales captured by cellular-level restrictions and axon caliber (Novikov et al., 2014; Fieremans et al., 2016; Lee et al., 2020b,c; Novikov, 2020). This behavior corresponds to short-range disorder in one dimension, yielding axial diffusivity time-dependence, given by (Appendix B.1)

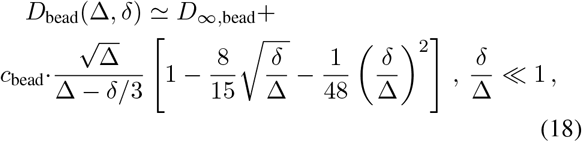

where the subscript “bead” denotes the caliber variation (beading). The above asymptotic behaviour in *δ*/Δ ≪ 1 limit actually works well for the wide-pulse case *δ*/Δ ∼ 1 with an error *<*1% in diffusivity time-dependence.

Recently, Abdollahzadeh et al. (2023) considered narrow axons with varying cross sections *A*(*z*) along the length *z* of axon skeleton and derived the exact relation between the cell geometry and diffusion properties based on the Fick-Jacob’s equation:

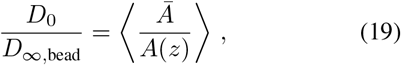

with *Ā* the mean cross-sectional area.

In simulations and axon diameter estimation using intra-axonal signals alone (Figure 5), the estimated axial diffusivity 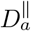 correlates with the long-time prediction *D*_∞,bead_ (19) due to beadings for axons with non-trivial caliber variations (Figure 7a), whereas this correlation is slightly weakened by the Rician noise at SNR = 100 (Figure 7b).

### 4.3. Variation of local axial and radial diffusivities

Simulations in real axons and fibers mimicking realistic axons show that axon size is underestimated due to caliber variations. This finding is unexpected and cannot be explained by current axonal diameter mapping models. Here we suggest accounting for this observation by introducing the variation of local axial and radial diffusivities. It is challenging to consider the variation of local axial and radial diffusivities at the same time, let alone the local axial kurtosis (see the full functional form in Appendix C). Therefore, we discuss the two cases individually with an assumption of using specific distributions for local diffusivity and decide which case better explains the observation. For simplicity, we choose to use a Gamma distribution to explain the variation of local diffusivity *D* (Jian et al., 2007),

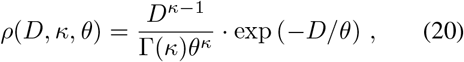

where Γ(·) is Euler’s Gamma function, and *κ* describes the shape and *θ* describes the scale of the distribution. The mean and variance of diffusivity *D* are *κθ* and *κθ*^2^, respectively.

#### 4.3.1. Variation of local radial diffusivity

In this case, we assume that local radial diffusivity varies along axons, whereas local axial diffusivity does not. The distribution of local radial diffusivity is described by a Gamma distribution 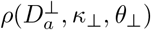 in Eq. (20). Given that the normalized spherical mean sig-nal *S*_*a*_(*b*) in an axonal segment is given by Eq. (7), the spherical mean signal summed over all axonal segments is

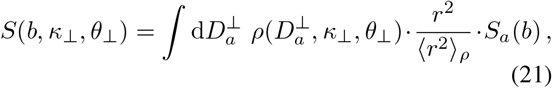

where 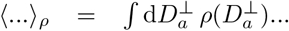, and the factor *r*^2^/ ⟨*r*^2^⟩_*ρ*_ accounts for the volume fraction variation of each axonal cross section. In the wide pulse limit, Neuman’s solution (9) suggests 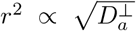. Substituting into Eq. (21), we obtain the spherical mean signal summed over all axonal segments,

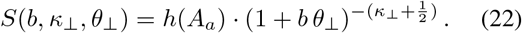

We fit the above equation to the simulated spherical mean signal in real axons (Figure 8a) and obtain *κ*_⊥_ *<* 0.001, *θ*_⊥_ = 0.96 *μ*m^2^/ms, and 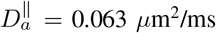, yielding an estimate of radius *<* 0.001 *μ*m. The estimated axial diffusivity and axon radius are both under-estimated with low quality-of-fit. Therefore, the variation of radial diffusivity fails to explain the observed underestimation of axon size.

**Figure 8:**
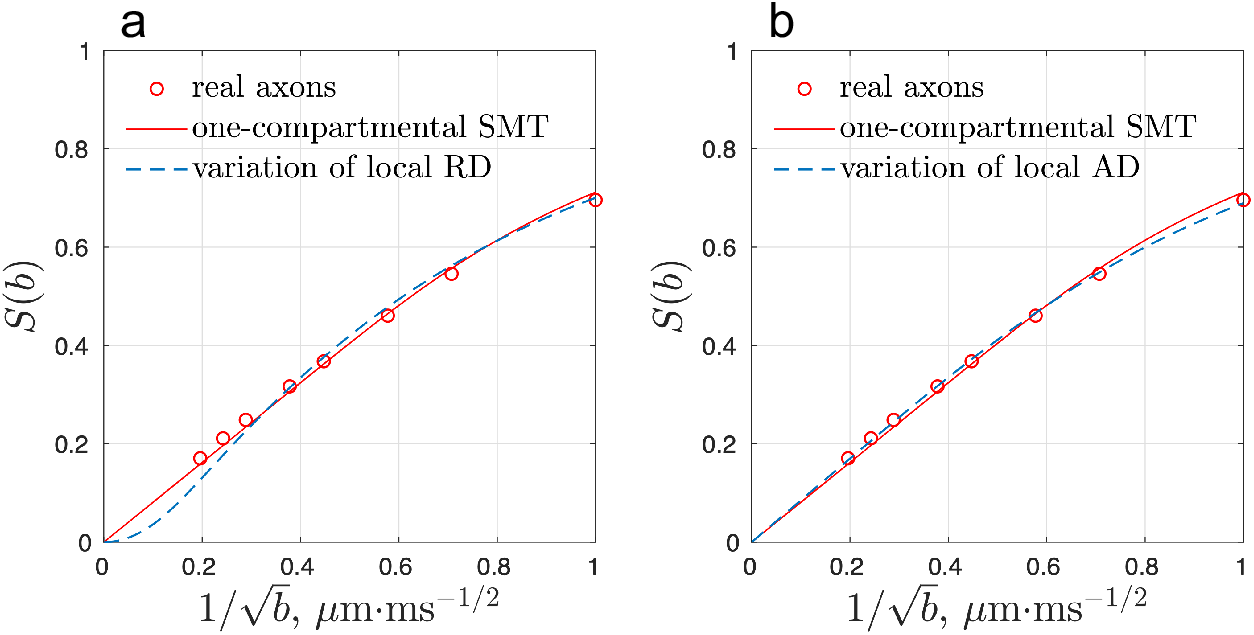
To explain the underestimated axon size due to caliber variations in Figure 5, we suggest accounting for this observation by using a Gamma distribution for variations of local diffusivity (a) transverse to or (b) parallel to axons in the intra-axonal signals (7). RD = radial diffusivity 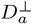, AD = axial diffusivity 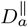.

#### 4.3.2. Variation of local axial diffusivity

In this case, we assume that the local axial diffusivity varies along the length of the axons, and local radial diffusivity is kept constant along the axons. The distribution of local axial diffusivity is described by a Gamma distribution 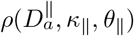. Similarly, the spherical mean signal summed over all axonal segments is

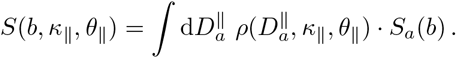

Substituting Eqs. (7) and (20) into the above equation, we obtain

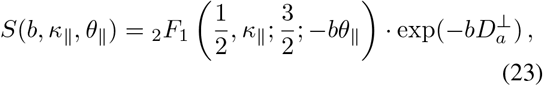

where _2_*F*_1_ is a Gauss hypergeometric function. We fit the above functional form to the simulated spherical mean signals in real axons (Figure 8b) and obtain *κ*_∥_ = 2.59, *θ*_∥_ = 0.58 *μ*m^2^/ms, and an estimated radius *<* 0.001 *μ*m. This corresponds an estimate of 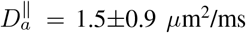. Though the estimated axon size is much smaller than the histological values (Figure 2a), the quality-of-fit for Gamma-distributed 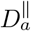 model (23) is slightly better that of the ordinary single-compartment SMT model (7). The estimated mean and standard deviation of local axial diffusivity may thus serve as biomarkers that reflect properties of the under-lying axonal beadings, as suggested in previous studies (Budde and Frank, 2010; Lee et al., 2020b).

### 4.4. Multi-compartmental AxCaliber-SMT

In white matter, the diffusion signals receive contributions from multiple components, including intracellular water, extra-cellular water, and CSF. To estimate axon size using biophysical modeling of the dMRI signal, we consider signal contributions from all compartments for the full range of b-values (Fan et al., 2020). To validate axonal diameter mapping models that account for multiple tissue compartments, we fit the multi-compartmental AxCaliber-SMT model to the volume-weighted sum of simulated intra-axonal signals and anisotropic (extra-axonal space) and isotropic (CSF) Gaussian signals. Our simulations show that caliber variations lead to underestimated axon size, whereas undulations lead to overestimated axon size. These findings are similar to the results of the simulations and model fitting for the intra-axonal signals in isolation, as captured by the single-compartment SMT model. Considering the contribution of signals from multiple compartments does not significantly alter the results of axon size estimation.

Simulations with added Rician noise (SNR = 100) show that the axon size is underestimated for axons with either strong caliber variations or strong undulations. This observation can be explained by the resolution limit in Section 4.1 (Nilsson et al., 2017; Andersson et al., 2022). It is worthwhile to note that the intra-axonal volume fraction *f*_*a*_ varies around 0.75 (0.5-1) in simulations of multiple-compartments, effectively resulting in a smaller SNR for intra-axonal signals. Therefore, in simulations of multiple compartments (Figure 6b), the bias due to noise is larger than that of the single-compartment SMT results (Figure 5b).

### 4.5. Interpretation of alterations in axon size estimates

Axonal diameter mapping using the spherical mean dMRI signal is affected by the Rician noise floor and axonal morphology, including caliber variations and undulations. Our findings from systematic simulations in axonal substrates with varying caliber and undulations suggest that the interpretation of alterations in MR-estimated axon size should be interpreted with caution, especially in pathological tissues. For instance, the observation of decreased axon size could be the result of axonal beading and/or noise, whereas the observation of increased axon size could be the result of undulations.

The sensitivity of dMRI signals to caliber variations and undulations has be demonstrated in (Budde and Frank, 2010; Brabec et al., 2020) and this study. Potential applications of these findings include monitoring axonal pathology, such as characterizing axonal undulations observed in human post-mortem brain tissue acutely following traumatic brain injury (TBI) (Tang-Schomer et al., 2012), and axonal varicosity (beading) observed following TBI (Johnson et al., 2013; Tang-Schomer et al., 2012) and ischemia in white matter (Garthwaite et al., 1999). Our findings suggest that axonal diameter mapping is potentially sensitive to axonal alterations, such as undulations in TBI patients and axonal beading in TBI and ischemic stroke patients.

### 4.6. Limitations

The ex vivo sample was fixed by immersion in paraformaldehyde and glutaraldehyde solution (Shapson-Coe et al., 2021). Undulations and caliber variations in the fixed tissue scanned with a serial sectioning EM may be stronger than in vivo potentially due to fixation time for large sample, shrinkage during tissue preparation, or even imperfect mechanical sectioning.

Furthermore, the extra-axonal space signals are generated based on a time-independent Gaussian ellipsoid in our simulations. However, diffusion in the extra-axonal space is non-Gaussian; the diffusivity and kurtosis time-dependence in the extra-axonal space is nontrivial, and may contribute to biases in the estimation of axon diameter if the signal is dominated by low b-values (Burcaw et al., 2015; Fieremans et al., 2016; Lee et al., 2018; De Santis et al., 2016). Extra-cellular space maintenance strategies for immersion fixation (Karlu-pia et al., 2023) could be applied to preserve and image the extra-cellular space using EM in the future study. Further segmentation and diffusion simulation in extracellular space would help to understand its contribution to the time-dependence at low b-values.

Finally, in this work, we have only considered axonal diameter mapping using the conventional pulsed-gradient spin-echo sequence, which is a linear tensor encoding scheme (Stejskal and Tanner, 1965). The effect of cellular-level features on axonal diameter mapping using other diffusion gradient waveforms (Eriksson et al., 2015; Westin et al., 2016), such as planar and spherical tensor encoding waveforms, is not considered in this study. In Appendix D, we show that the resolution limit (smallest detectable axon size) of planar and spherical tensor encoding waveform is slightly smaller (i.e., better) than that of linear tensor encoding for a given b-value *<* 10 ms/*μ*m^2^ (planar tensor encoding) and *<* 5 ms/*μ*m^2^ (spherical tensor encoding). However, due to limitations in slew rate and echo time, generalized diffusion gradient waveform aside from linear tensor encoding generally achieve much lower b-values than linear tensor encoding. It is difficult to distinguish intra-axonal and extra-axonal signals at low b-values, and thus using generalized waveforms is still less efficient than using linear tensor encoding for axonal diameter mapping.

## 5. Conclusions

Monte Carlo simulations of diffusion in realistic axonal substrates and their fiber derivatives show that axonal diameter mapping in white matter axons is affected by caliber variations and undulations, the two salient axonal features in fixed human brain tissue imaged by EM. Applying axonal diameter mapping on the simulated spherical mean signal demonstrate underestimated axon size in axons with strong caliber variations and overestimated axon size in those with strong undulations. This finding suggests that the interpretation of alterations in MR-estimated axon caliber in studies of pathological white matter tissue should factor in caliber variations and undulations as potential contributors to observed decreases or increases in axon size, respectively. The relevance of these finding to in vivo axonal diameter mapping will require further exploration and systematic multi-modality validation.

## Acknowledgements

We would like to thank Matthew P. Frosch, Daniel R. Berger, and Jeff W. Lichtman for the fruitful discussion. The research reported in this manuscript is supported by the Office of The Director (OD) of the National Institutes of Health (NIH) and National Institute of Dental & Craniofacial Research (NIDCR) under the award number DP5 OD031854, National Institute of Aging of the NIH under the award number K99 AG073506, and the National Institute of Neurological Disorders and Stroke of the NIH under award numbers R01 NS118187, R21 NS081230 and R01 NS088040, and by the National Institute of Biomedical Imaging and Bioengineering (NIBIB) of the NIH under award numbers U01 EB026996, P41 EB015896, P41 EB030006 and P41 EB017183.

Figure 1a is adapted from (Shapson-Coe et al., 2021) with permission from bioRxiv.

## Appendix A Axial diffusivity time-dependence due to undulations

For simplicity, we provide an approximate solution of axial diffusivity time-dependence in a single harmonic undulating fiber with no caliber variations, perturbatively in *w*_0_/*λ* ≪ 1, cf. *w*_0_/*λ* ≈ 0.11 in real axons (Figure 2b). In the previous study (Lee et al., 2020a), we show that, in narrow pulse limit, the axial diffusivity along a undulating fiber at diffusion time *t* is given by

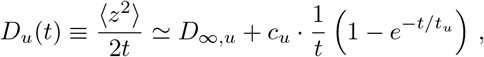

where long-time diffusivity *D*_∞,*u*_, strength of diffusivity time-dependence *c*_*u*_, and correlation time *t*_*u*_ are defined as follows:

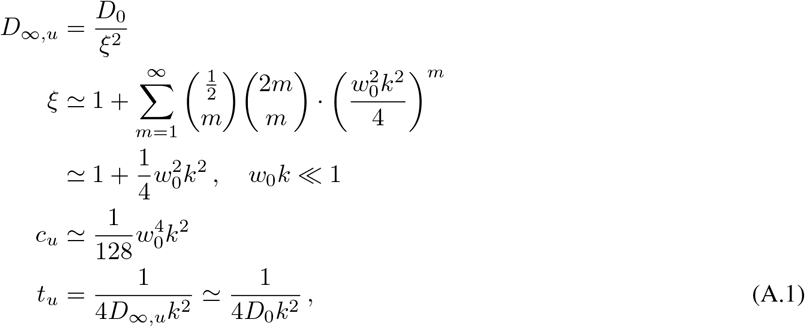

with undulation amplitude *w*_0_ and wavelength *λ* (*k* = 2*π*/*λ*). The approximation *D*_∞,*u*_ ≃ *D*_0_ in Eq. (A.1), i.e., *ξ* ≳ 1, is supported by the small value of *w*_0_*k* ∼ *w*_0_/*λ* in real axons (Figure 2b).

Then it is straightforward to calculate the instantaneous diffusivity in *d* = 1-dimension along the fiber,

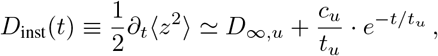

which yields the frequency *ω*-dependent dispersive diffusivity in a standard way (Burcaw et al., 2015; Novikov et al., 2019)

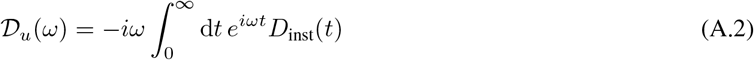

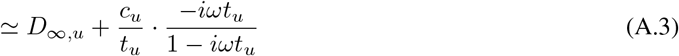

The knowledge of dispersive diffusivity enables to evaluate the diffusion signal *S* up to the second order cumulant in any sequence:

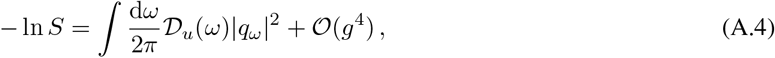

where *q*_*ω*_ is the Fourier transform of the diffusion wave vector 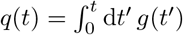. For a pulsed-gradient sequence of inter-pulse time interval Δ and pulse width *δ*, we have (Callaghan, 1993; Burcaw et al., 2015)

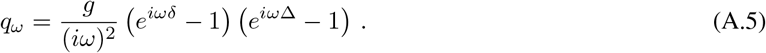

Substituting Eqs. (A.3) and (A.5) into Eq. (A.4), we obtain the axial diffusivity 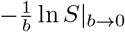 measured by the wide pulsed-gradient sequence due to undulations:

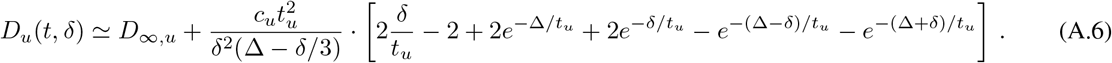

It is interesting that the diffusivity time-dependence transverse and parallel to a undulating fiber (with no caliber variations) shows the same functional form, with different scales in long-time diffusivity (0 vs *D*_∞,*u*_), strength of diffusivity time-dependence 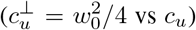, and correlation time 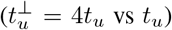 (Lee et al., 2020a). In wide pulse limit of undulations, i.e., *δ* ≫ *t*_*u*_, the axial diffusivity due to undulations in Eq. (A.6) acquires the Neuman (1974) form in Eq. (16).

## Appendix B Axial diffusivity time-dependence due to caliber variations

Details in tissue micro-geometry are homogenized by diffusion at long time, resulting in a coarse-grained effective medium with local varying diffusivity (Novikov and Kiselev, 2010; Novikov et al., 2014), manifested by the powerlaw scaling in time-dependent instantaneous diffusivity (Novikov et al., 2014):

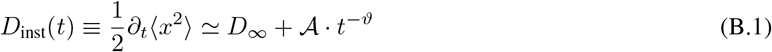

with long-time diffusivity *D*_∞_, strength of restrictions 𝒜, and dynamical exponent *ϑ* = (*p* + *d*)/2. The long-range density fluctuations of microstructure in *d*-dimension is described by the structural exponent *p* via its power spectrum. In narrow pulse limit, the typical cumulant diffusion coefficient

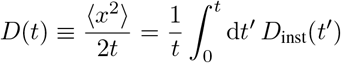

shows the same power-law scaling

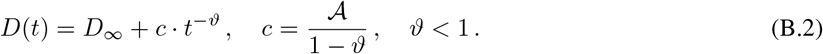

Substituting Eq. (B.1) into Eq. (A.2) yields the frequency *ω*-dependent dispersive diffusivity (Novikov and Kiselev, 2010; Novikov et al., 2014; Lee et al., 2020c):

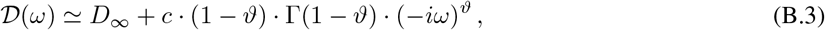

where Γ(·) is Euler’s Γ-function. Further, substituting Eqs. (B.3) and (A.5) into Eq. (A.4), we obtain the general form of time-dependent diffusivity 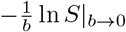 measured by using wide pulsed-gradient sequence:

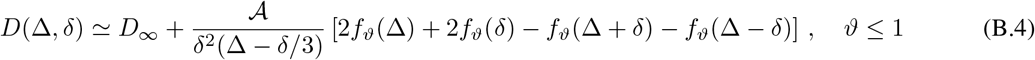

where

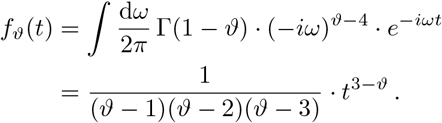

The relation Γ(1 − *ϑ*) · Γ(*ϑ*) = *π*/ sin(*πϑ*) is applied to simplify the equation. We will use this generalized functional form to discuss the time-dependent axial diffusivity due to caliber variations.

### Appendix B.1. Short-range disorder in 1d: AD time-dependence due to caliber variations

Diffusion along neurites is hindered by caliber variations, such as beadings and spines, whose random arrangement along neurites in histology (Hellwig et al., 1994; Glantz and Lewis, 2000; Morales et al., 2014) indicates the short-range disorder in 1-dimension (*p* = 0, *d* = 1). This corresponds to a dynamical exponent *ϑ* = 1/2 and the diffusivity measured by using wide pulsed-gradient sequence in Eq. (B.4):

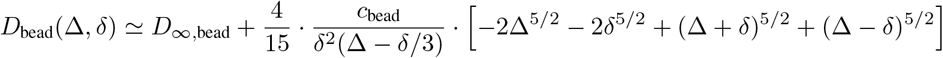

where *c*_bead_ = 2A is substituted based on Eq. (B.2), and the subscript “bead” denotes beadings. The asymptotic solution for narrow pulse, i.e., *δ*/Δ ≪ 1, is shown in Eq. (18).

## Appendix C The effect of local axial diffusivity variation along individual axon on axonal diameter mapping

In Eq. (7), axonal segments are assumed to be in cylindrical shape with the same local axial diffusivity 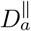 along each segment. However, the undulations and caliber variations along individual axon lead to variations of local axial diffusivity in each axonal segment, i.e., 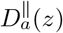 varying through axon’s main axis *z*. In addition, caliber variations *r*(*z*) lead to variations of local radial diffusivity 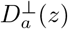 in each axonal segment as well. The former effect was not considered in previous studies, and the latter one was approximated by the first order expansion (Veraart et al., 2020). To account for these effects, the variation of local axial diffusivity and radial diffusivity in each axonal segment is considered in the calculation of spherical mean signal:

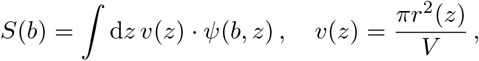

where d*z v*(*z*) and *V* = ∫ d*z v*(*z*) are volumes of an axonal segment at *z* and the whole axon, respectively. The spherical mean signal *ψ* (*b, z*) of an axonal segment has the same functional form as in Eq. (7), except that 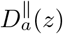 and 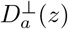 both vary in each segment. To estimate local diffusivity, 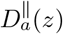 and 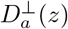, along and transverse to axonal segments, it is necessary to perform MC simulations of diffusion with random walkers initialized in each axonal segment at *z*. This is very time-consuming to perform such simulations with reasonable precision of local diffusivity estimations.

In addition to local diffusivity, the variation of local axial kurtosis 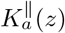 along individual axon yields the next order correction (Veraart et al., 2019):

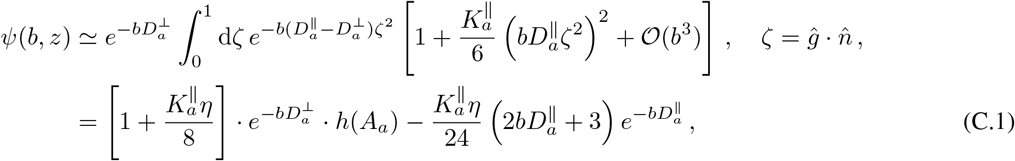

where *ĝ* and 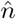 are directions of diffusion gradient and axon segment, and

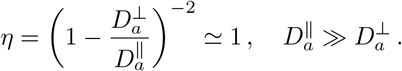

The second right-hand-side term in Eq. (C.1) decays much faster than the first term due to the exponential decay 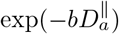. Similarly, the local diffusivities 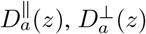 and local axial kurtosis 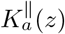 have to be estimated by performing diffusion simulations with random walkers initialized in each axonal segment located at *z*.

## Appendix D Resolution limit for axonal diameter mapping using generalized diffusion gradient waveform

For the generalized diffusion gradient waveform, the q-space trajectory imaging (Westin et al., 2016) can be described by a general B-tensor. The diffusion weighting tensor is given by

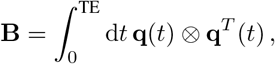

where 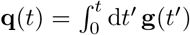, and Larmor gradient **g**(*t*) is the product of diffusion gradient and gyromagnetic ratio. For an axi-symmetric B-tensor, its axial and radial components, *b*_∥_ and *b*_⊥_, define its b-value (trace) *b* = *b*_∥_ + 2*b*_⊥_ and anisotropy 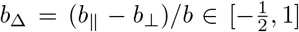. For conventional LTE, *b*_Δ_ = 1; for STE, *b*_Δ_ = 0; for PTE, 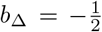. Considering an axi-symmetric signal kernel (i.e., cylinder in this case), the spherical mean signal measured by the axi-symmetric B-tensor waveform is given by (Eriksson et al., 2015)

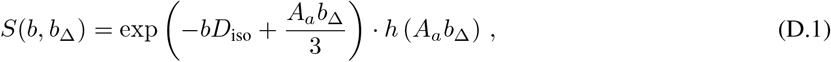

Where 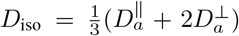, and *h*(·) and *A*_*a*_ are defined in Eqs. (6) and (8). In the low frequency limit, the frequency *ω*-dependent dispersive diffusivity for resticted diffusion is Re𝒟(*ω*) ∼ *ω*^2^, and the radial diffusivity of a straight cylinder for an arbitrary gradient waveform can be approximated by (Burcaw et al., 2015; Nilsson et al., 2017; Novikov et al., 2019)

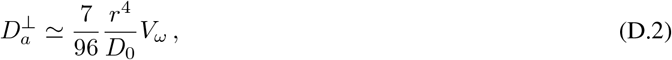

where *V*_*ω*_ is the spectral encoding variance defined from the second order of **q**(*ω*) and is independent of the scaling of gradient strength:

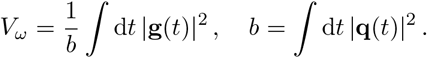

In practice, the frequency spectrum of a general B-tensor waveform could be different in *q*_*x*_, *q*_*y*_, and *q*_*z*_ components. In Eq. (D.1), the 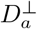 is assumed to be similar when it is measured by *q*_*x*_, *q*_*y*_, *q*_*z*_, and their vector combinations. Furthermore, the relation (D.2) between 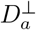 and *r* is approximated via the average of 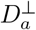 measured by applying *q*_*x*_, *q*_*y*_, and *q*_*z*_, respectively. We will use the above relations to evaluate the resolution limit for axonal diameter mapping using axi-symmetric B-tensor encoding sequences.

### Appendix D.1. Resolution limit for axonal diameter mapping using STE and LTE spherical mean signal

To evaluate the resolution limit of axonal diameter mapping using B-tensor encoding sequence with *b*_Δ_ ∈ [0, 1], we calculate the difference of spherical mean signals (D.1) between infinitely thin cylinders and those with finite radius, cf. Eq. (11):

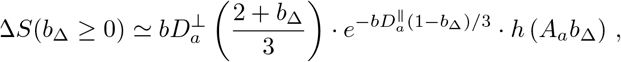

where we approximate 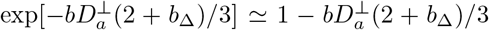. Substituting the Δ*S*(*b*_Δ_ ≥ 0) and 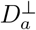 in Eq. (D.2) into the requirement (13) of resolution limit for axonal diameter mapping, we obtain

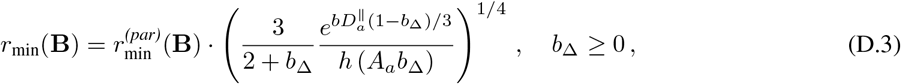

where the 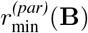 is analogous to the resolution limit for axonal diameter mapping by applying generalized LTE waveform (not just pulsed-gradient) of the same *V*_*ω*_ value transverse to highly aligned cylinders (Nilsson et al., 2017):

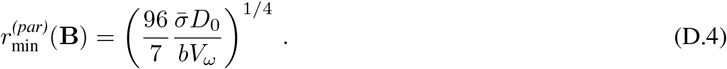

Unlike LTE, non-LTE waveform has the additional signal sensitivity to diffusion parallel to cylinders, even when the main axis of B-tensor waveform is transverse to cylinders. This is manifested by the fact that *r*_min_(**B**) is not the same as 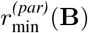 for STE waveform (*b*_Δ_ = 0):

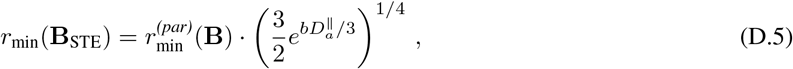

where *h*(*x*) = 1 for *x* → 0^+^ is applied.

For LTE waveform (*b*_Δ_ = 1), the resolution limit is given by

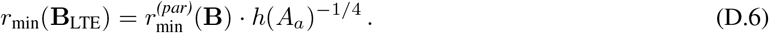

Substituting *V*_*ω*_ = 2/*δ*(Δ − *δ*/3) for the pulsed-gradient LTE into the above equation yields Eq. (14).

### Appendix D.2. Resolution limit for axonal diameter mapping using PTE spherical mean signal

To evaluate the resolution limit of axonal diameter mapping using B-tensor encoding sequence with 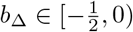, we calculate the difference of spherical mean signals (D.1) between infinitely thin cylinders and those with finite radius, c.f., Eq. (11):

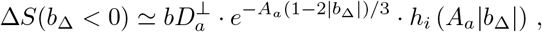

where we approximate 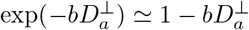, and

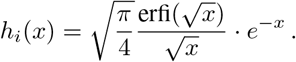

Substituting the Δ*S*(*b*_Δ_ *<* 0) and 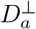 in Eq. (D.2) into the requirement (13) of resolution limit for axonal diameter mapping, we have

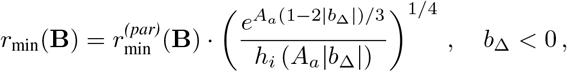

For PTE waveform 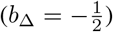, the resolution limit for axonal diameter mapping is

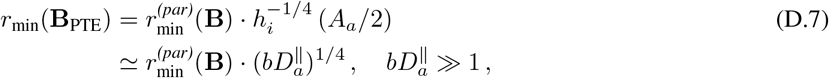

where *h*_*i*_(*x*) ≃ 1/(2*x*) for *x* ≫1. Substituting Eq. (D.4) into the above equation, we notice that the resolution limit for PTE waveform is independent of *b* at high b-values,

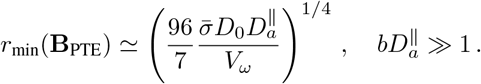

In other words, at high b-values, the resolution limit for PTE waveform will not be significantly improved by applying stronger gradients.

We compare the resolution limit for axonal diameter mapping using LTE (D.6), PTE (D.7), and STE (D.5) within total time = 30 ms for diffusion gradient application. The LTE waveform is initialized with a pulsed-gradient sequence of an inter-pulse duration Δ = 20 ms and a pulse width *δ* = 10 ms. The LTE, PTE, and STE are optimized and Maxwell-compensated based on a b-value maximizing framework (Szczepankiewicz et al., 2019), where we set the maximal gradient strength and slew rate at 500 mT/m and 600 T/m/s to create the waveform, consistent with the spec of Connectom 2.0 scanner (Huang et al., 2021). Then we scale the diffusion gradient amplitude for a given b-value without applying constraints on maximal gradient strength and slew rate. The resolution limit is calculated for SNR = 100, *n* = 60, and *z*_*α*_ = 1.64 for *α* = 0.05. The intrinsic diffusivity and axial diffusivity are fixed at *D*_0_ = 2 *μ*m^2^/ms and 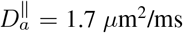. For a given b-value, PTE and STE provide slightly smaller (i.e., better) resolution limit than LTE for b-value *<* 10 ms/*μ*m^2^ and *<* 5 ms/*μ*m^2^, respectively (Figure D.1). However, for a given maximal gradient strength (e.g., 300 mT/m for Connectom scanner 1.0 (Fan et al., 2022)), PTE and STE can only achieve much smaller b-values, where the intra- and extra-axonal signals are difficult to distinguish. Therefore, LTE is still the most efficient waveform for axonal diameter mapping so far.

**Figure D.1:**
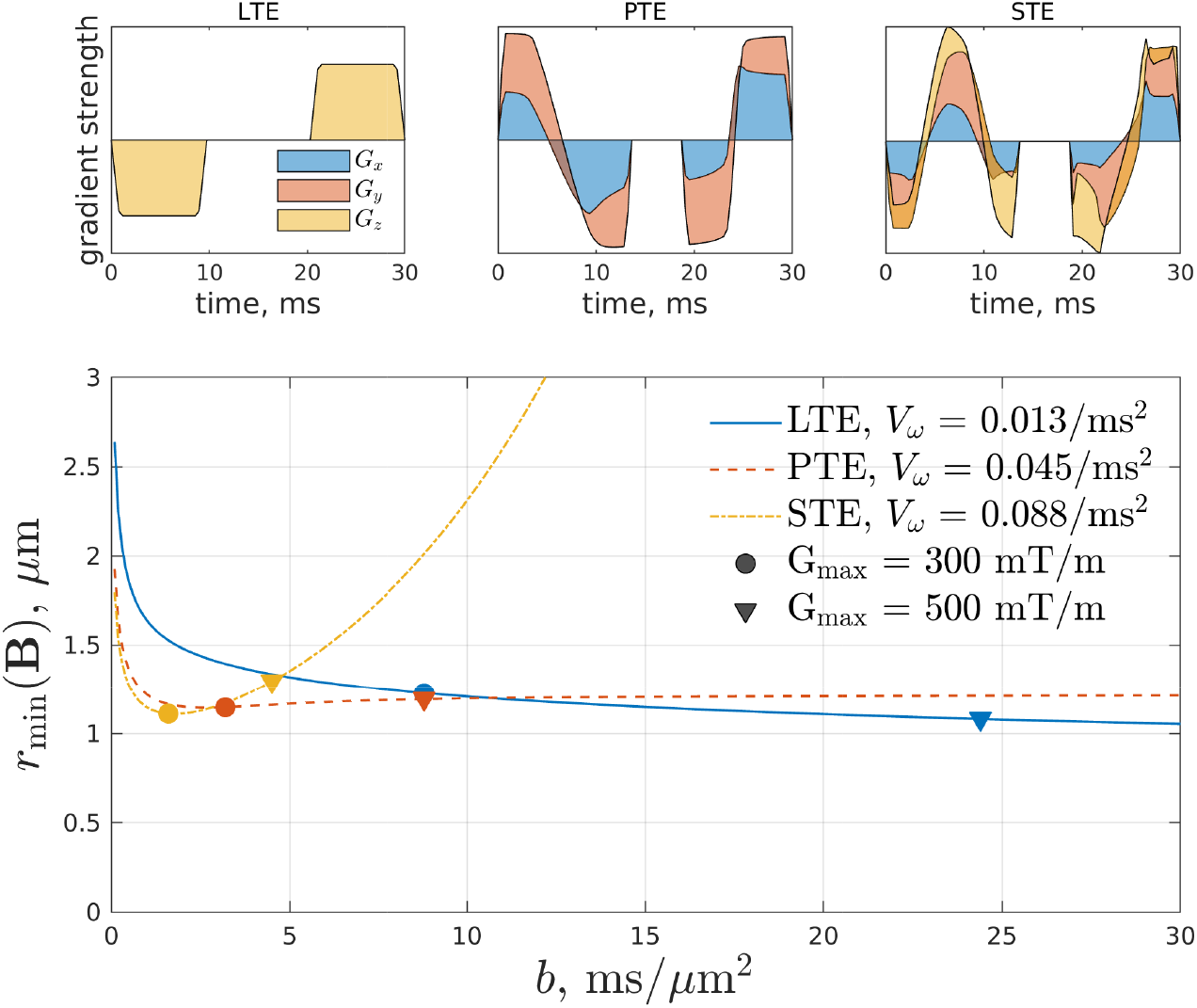
Resolution limit, i.e., smallest detectable axon size *r*_min_(**B**), for axonal diameter mapping using LTE (D.6), PTE (D.7), and STE (D.5) waveform. For a given b-value, PTE and STE lead to slightly smaller resolution limit than LTE for b-value *<* 10 ms/*μ*m^2^ and *<* 5 ms/*μ*m^2^, respectively. However, PTE and STE require much larger gradient strength to achieve the same b-values.

## Supporting Information

### S1. Power-law scaling at high b-value

At high *b*-value, the extra-axonal space and CSF signals decay away exponentially, while the deviation from the slow, 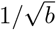 -scaling of the intra-axonal signals provide an estimation of axon diameter (Veraart et al., 2020). By fitting this unique power-law scaling to the combined intra-axonal, extra-axonal and CSF signals at high *b >* 6 ms/*μ*m^2^, we further test the effect of axonal geometry on ADM when the extra-axonal and CSF signal contributions are included (though negligible in modeling at high *b*).

#### S1.1. Methods

The single-compartment SMT model only applies to intra-axonal signals, whereas diffusion signals in white matter tissues have contributions from multiple compartments, including the extra-axonal space and CSF. To extend the applicability of axonal diameter mapping using the spherical mean signal to white matter, Veraart et al. (2020) proposed to fit the intra-axonal SMT model to diffusion signals at high b-values, e.g, *>* 6 ms/*μ*m^2^ in vivo, in the regime where the diffusion signals in the extra-axonal space and CSF decay exponentially and become negligible. At high b-values, the dominant contribution to the spherical mean signal comes from the intra-axonal space:

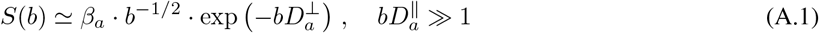

where

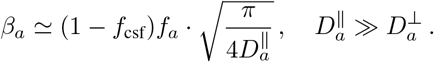

For the above functional form, in Eq. (6) we approximate 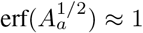 for 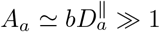 at high b-value. Hence, we have a two-parameter fit with parameters *β*_*a*_ and 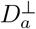. The estimated 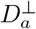 is then translated into the MR-estimated axon size using Eq. (9).

To evaluate ADM performance, the multi-compartmental model in Eq. (A.1) is fitted to the combined signals in Eq. (3) using nonlinear least squares with positive constraints for all parameters. The magnitude signals are composed of MC-simulated signals in the intra-axonal space and added signals in the extra-axonal space and CSF with and without Rician noise, where the noise levels in the real and imaginary parts were both *σ* = *S*_0_/SNR with *S*_0_ ≡ 1 and SNR = ∞ (no noise) and 100, respectively.

#### S1.2. Results

In white matter voxels, diffusion signals receive contributions from multiple components, including the intra-axonal space, extra-axonal space, and CSF. To validate axonal diameter mapping in white matter, we combine the intraaxonal MC-simulated signals with axi-symmetric Gaussian signals representing the extra-axonal space and isotropic Gaussian signals representing CSF. We fit the power-law scaling model in Eq. (A.1) to the combined signals in Eq. (3) at high b-values *>* 6 ms/*μ*m^2^. The noiseless combined signals lead to overestimated axon size in axons with strong undulations, and underestimated axon size in axons with strong caliber variations (Figure S4a). However, the combined signals with added Rician noise (SNR = 100) result in underestimated axon size due to the Rician noise floor, for both cases of undulations and caliber variations (Figure S4b).

**Figure S1:**
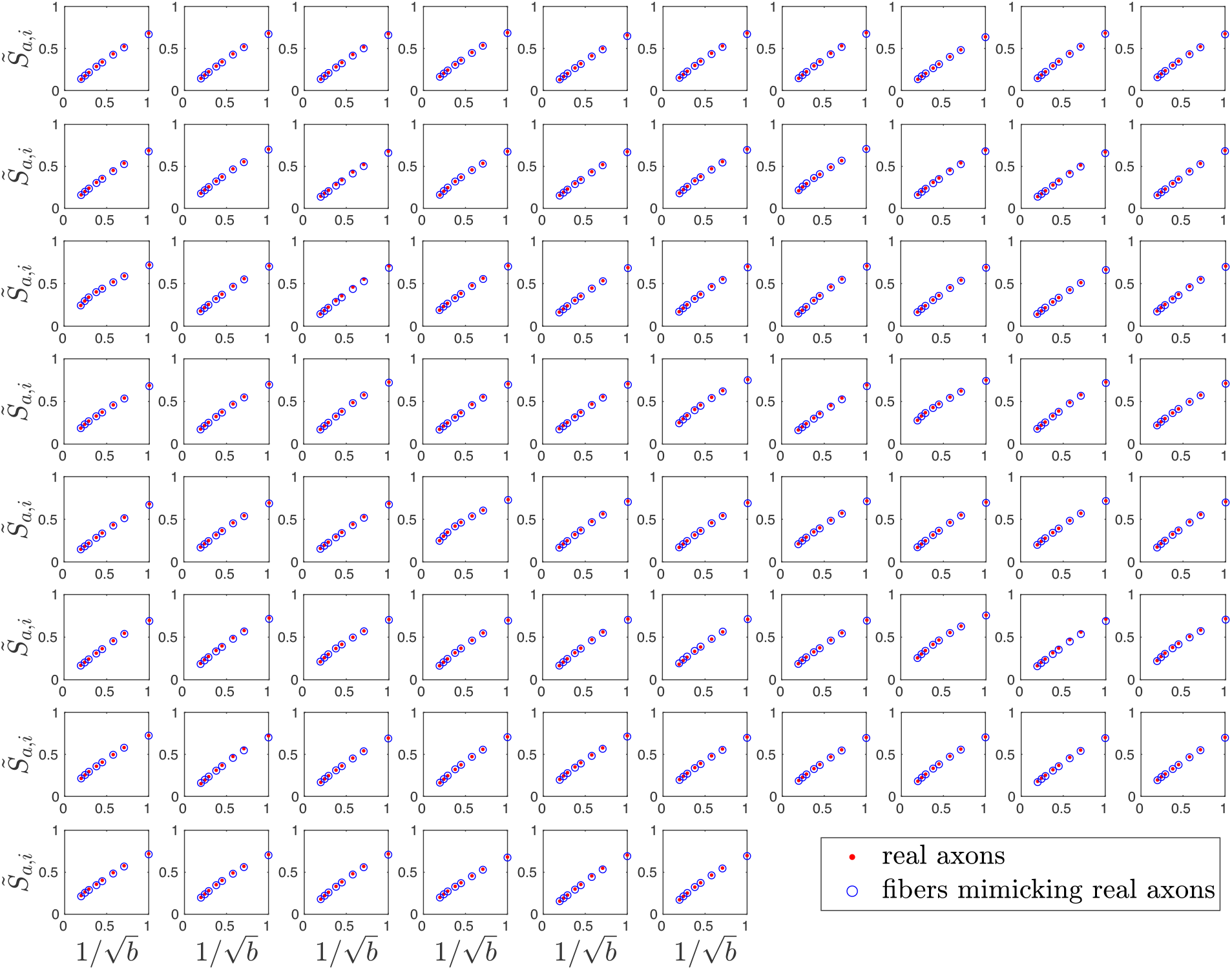
Simulated spherical mean signals in 76 undulating, beaded fibers of circular cross sections (blue) with the same caliber variations and undulations as in real axons (red), i.e., 100% CV(*r*) and 100% *w*_0_ in Figure 3. Simulation results in real axons coincide with those in corresponding fiber derivatives of the same features.

**Figure S2:**
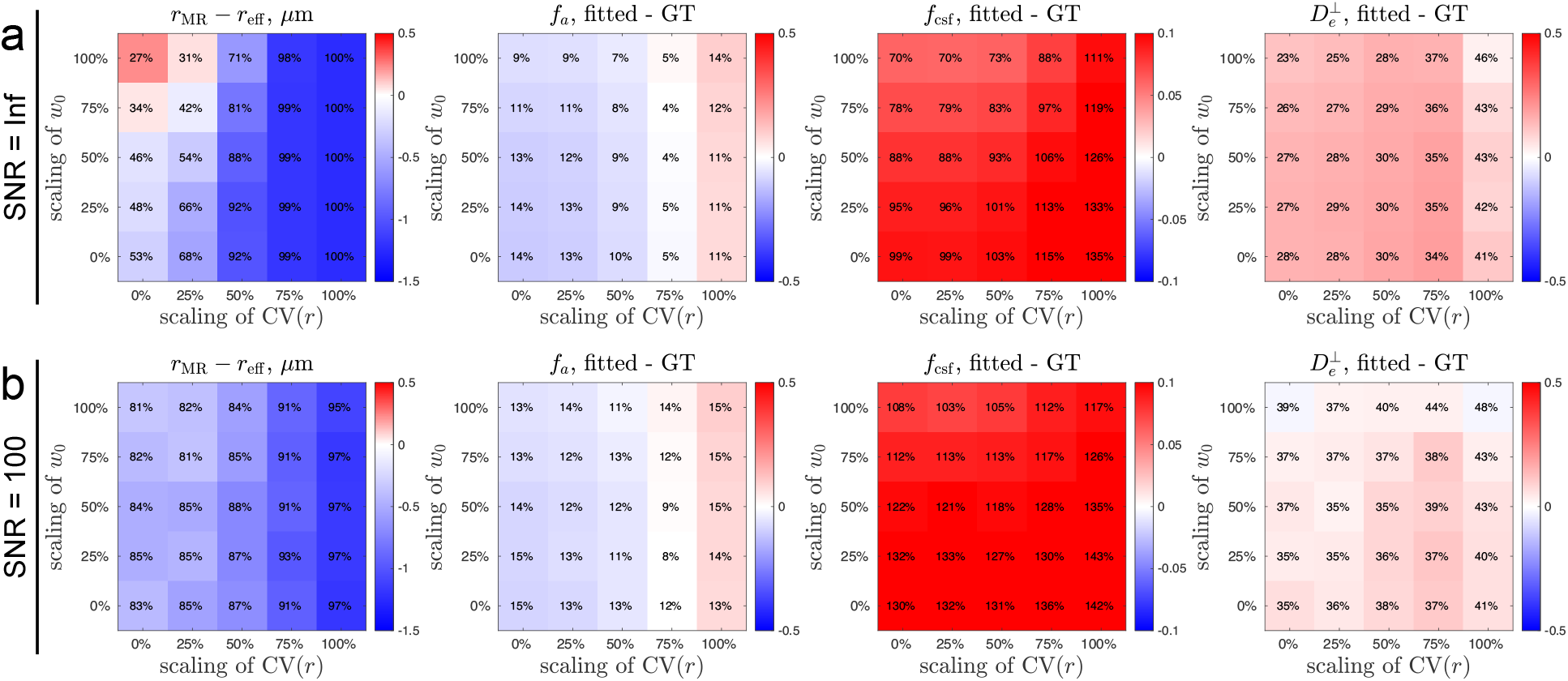
Axon size estimation based on diffusion simulations in undulating, beaded fibers reveals the effect of undulations and caliber variations on axonal diameter mapping, respectively. The multi-compartmental AxCaliber-SMT model 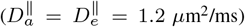 in Section 2.6.2 is fitted to the multi-compartmental signal in Section 2.5.2. The number in each pixel indicates the NRMSE between the estimation and ground truth. When a parameter is underestimated, its NRMSE cannot exceed 100% due to the positive constraint in nonlinear least square fitting. (a) Simulations without adding the noise show that strong undulations lead to slight overestimation of axon size, and caliber variations lead to underestimation of axon size. In contrast, the estimated intra-axonal volume fraction *f*_*a*_ shows the opposite trend; undulations lead to slight underestimation of *f*_*a*_, and caliber variations lead to overestimation of *f*_*a*_. In addition, both undulations and caliber variations lead to overestimation of CSF volume fraction *f*_csf_ and extra-axonal radial diffusivity 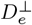. (b) Simulations with Rician noise (SNR = 100) show that the Rician noise floor leads to underestimation of axon size with low precision, manifested by large NRMSE. The bias in *f*_*a*_, *f*_csf_, and 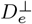 is similar to the noiseless result, and yet the precision is lower, i.e., larger NRMSE. The radial diffusivity 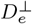 of the extra-axonal space is in units of *μ*m^2^/ms. GT = ground truth.

**Figure S3:**
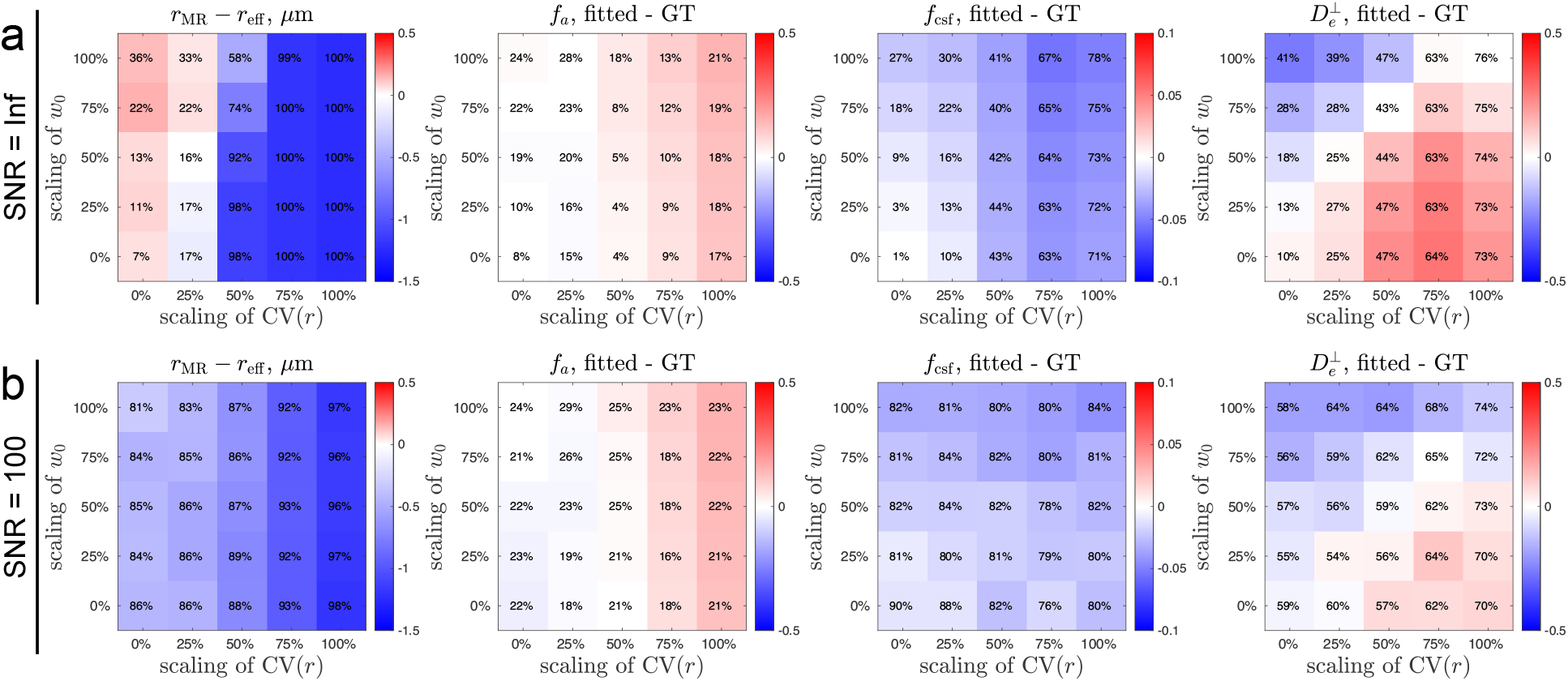
Axon size estimation based on diffusion simulations in undulating, beaded fibers reveals the effect of undulations and caliber variations on axonal diameter mapping, respectively. The multi-compartmental AxCaliber-SMT model 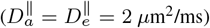 in Section 2.6.2 is fitted to the multi-compartmental signal in Section 2.5.2. The number in each pixel indicates the NRMSE between the estimation and ground truth. When a parameter is underestimated, its NRMSE cannot exceed 100% due to the positive constraint in nonlinear least square fitting. (a) Simulations without adding the noise show that undulations lead to overestimation of axon size, and caliber variations lead to underestimation of axon size. Further, undulations do not lead to significant bias in intra-cellular volume fraction *f*_*a*_, and caliber variations lead to overestimation of *f*_*a*_. In addition, both undulations and caliber variations lead to underestimation of CSF volume fraction *f*_csf_. Finally, undulations lead to underestimation of extra-axonal radial diffusivity 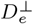, and yet caliber variations lead to overestimation of 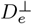. (b) Simulations with Rician noise (SNR = 100) show that the Rician noise floor leads to underestimation of axon size with low precision, manifested by large NRMSE. The bias in *f*_*a*_, *f*_csf_, and 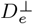 is similar to the noiseless result, and yet the precision is lower, i.e., larger NRMSE. The radial diffusivity 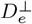 of the extra-axonal space is in units of *μ*m^2^/ms. GT = ground truth.

**Figure S4:**
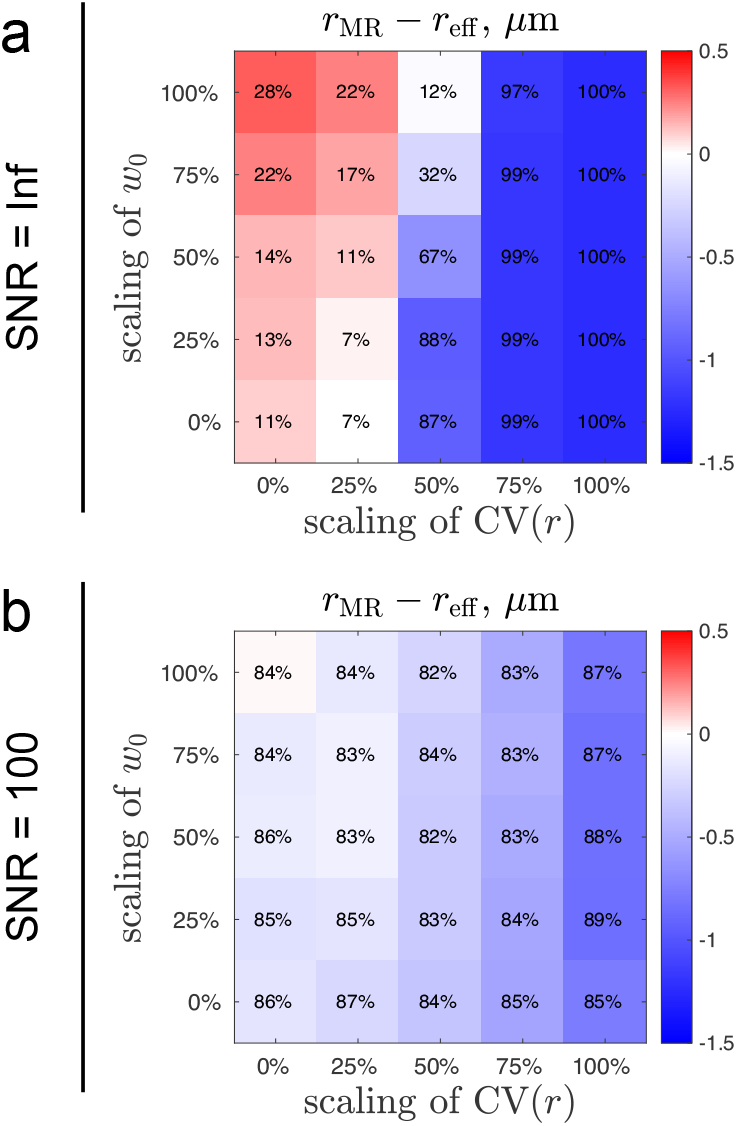
Axon size estimation based on diffusion simulations in undulating, beaded fibers reveals the effect of undulations and caliber variations on axonal diameter mapping, respectively. The power-law scaling model in Section S1 is fitted to the multi-compartmental signal at high b-values (*b >* 6 ms/*μ*m^2^) in Section 2.5.2. The number in each pixel indicates the NRMSE between the estimation *r*_MR_ and histology *r*_eff_. When a parameter is underestimated, its NRMSE cannot exceed 100% due to the positive constraint in nonlinear least square fitting. (a) Simulations without adding the noise show that undulations lead to overestimation of axon size, and caliber variations lead to underestimation of axon size. (b) Simulations with Rician noise (SNR = 100) show that Rician noise floor leads to underestimation of axon size with low precision, manifested by large NRMSE.

## References

Abdollahzadeh, A., Belevich, I., Jokitalo, E., Tohka, J., Sierra, A., 2019. Automated 3D axonal morphometry of white matter. Scientific reports 9, 6084.

Abdollahzadeh, A., Coronado-Leija, R., Mehrin, S., Lee, H.H., Sierra, A., Fieremans, E., Novikov, D.S., 2023. Characterization of white matter myelinated and unmyelinated axons from diffusion MRI perspective. 31st Annual Meeting of the International Society for Magnetic Resonance in Medicine, Toronto, ON, Canada.

Alexander, D.C., Hubbard, P.L., Hall, M.G., Moore, E.A., Ptito, M., Parker, G.J., Dyrby, T.B., 2010. Orientationally invariant indices of axon diameter and density from diffusion MRI. NeuroImage 52, 1374–1389.

Andersson, M., Pizzolato, M., Kjer, H.M., Skodborg, K.F., Lundell, H., Dyrby, T.B., 2022. Does powder averaging remove dispersion bias in diffusion mri diameter estimates within real 3d axonal architectures? NeuroImage 248, 118718.

Assaf, Y., Blumenfeld-Katzir, T., Yovel, Y., Basser, P.J., 2008. AxCaliber: a method for measuring axon diameter distribution from diffusion MRI. Magnetic Resonance in Medicine: An Official Journal of the International Society for Magnetic Resonance in Medicine 59, 1347–1354.

Brabec, J., Lasič, S., Nilsson, M., 2020. Time-dependent diffusion in undulating thin fibers: Impact on axon diameter estimation. NMR in Biomedicine 33, e4187.

Budde, M.D., Frank, J.A., 2010. Neurite beading is sufficient to decrease the apparent diffusion coefficient after ischemic stroke. Proceedings of the National Academy of Sciences 107, 14472–14477.

Burcaw, L.M., Fieremans, E., Novikov, D.S., 2015. Mesoscopic structure of neuronal tracts from time-dependent diffusion. NeuroImage 114, 18–37.

Callaghan, P., Jolley, K., Lelievre, J., 1979. Diffusion of water in the endosperm tissue of wheat grains as studied by pulsed field gradient nuclear magnetic resonance. Biophysical journal 28, 133–141.

Callaghan, P.T., 1993. Principles of nuclear magnetic resonance microscopy. Oxford University Press on Demand.

Cluskey, S., Ramsden, D., 2001. Mechanisms of neurodegeneration in amyotrophic lateral sclerosis. Molecular Pathology 54, 386.

De Santis, S., Jones, D.K., Roebroeck, A., 2016. Including diffusion time dependence in the extra-axonal space improves in vivo estimates of axonal diameter and density in human white matter. NeuroImage 130, 91–103.

DeLuca, G., Ebers, G., Esiri, M., 2004. Axonal loss in multiple sclerosis: a pathological survey of the corticospinal and sensory tracts. Brain 127, 1009–1018.

Deoni, S.C., Rutt, B.K., Arun, T., Pierpaoli, C., Jones, D.K., 2008. Gleaning multicomponent t1 and t2 information from steady-state imaging data. Magnetic Resonance in Medicine: An Official Journal of the International Society for Magnetic Resonance in Medicine 60, 1372–1387.

Dhital, B., Reisert, M., Kellner, E., Kiselev, V.G., 2019. Intra-axonal diffusivity in brain white matter. NeuroImage 189, 543–550.

Drobnjak, I., Zhang, H., Ianuş, A., Kaden, E., Alexander, D.C., 2016. Pgse, ogse, and sensitivity to axon diameter in diffusion mri: Insight from a simulation study. Magnetic Resonance in Medicine 75, 688–700.

Eriksson, S., Lasič, S., Nilsson, M., Westin, C.F., Topgaard, D., 2015. Nmr diffusion-encoding with axial symmetry and variable anisotropy: Distinguishing between prolate and oblate microscopic diffusion tensors with unknown orientation distribution. The Journal of chemical physics 142, 104201.

Fan, Q., Eichner, C., Afzali, M., Mueller, L., Tax, C.M., Davids, M., Mahmutovic, M., Keil, B., Bilgic, B., Setsompop, K., et al., 2022. Mapping the human connectome using diffusion mri at 300 mt/m gradient strength: Methodological advances and scientific impact. NeuroImage 254, 118958.

Fan, Q., Nummenmaa, A., Witzel, T., Ohringer, N., Tian, Q., Setsompop, K., Klawiter, E.C., Rosen, B.R., Wald, L.L., Huang, S.Y., 2020. Axon diameter index estimation independent of fiber orientation distribution using high-gradient diffusion mri. Neuroimage 222, 117197.

Fieremans, E., Burcaw, L.M., Lee, H.H., Lemberskiy, G., Veraart, J., Novikov, D.S., 2016. In vivo observation and biophysical interpretation of time-dependent diffusion in human white matter. NeuroImage 129, 414–427.

Fieremans, E., Lee, H.H., 2018. Physical and numerical phantoms for the validation of brain microstructural MRI: A cookbook. NeuroImage 182, 39–61.

Garthwaite, G., Brown, G., Batchelor, A., Goodwin, D., Garthwaite, J., 1999. Mechanisms of ischaemic damage to central white matter axons: a quantitative histological analysis using rat optic nerve. Neuroscience 94, 1219–1230.

van Gelderen, P., de Vleeschouwer, M.H., DesPres, D., Pekar, J., van Zijl, P.C., Moonen, C.T., 1994. Water diffusion and acute stroke. Magnetic Resonance in Medicine 31, 154–163.

Glantz, L.A., Lewis, D.A., 2000. Decreased dendritic spine density on prefrontal cortical pyramidal neurons in schizophrenia. Archives of general psychiatry 57, 65–73.

Grussu, F., Schneider, T., Yates, R.L., Zhang, H., Wheeler-Kingshott, C.A.G., DeLuca, G.C., Alexander, D.C., 2016. A framework for optimal whole-sample histological quantification of neurite orientation dispersion in the human spinal cord. Journal of neuroscience methods 273, 20–32.

Gudbjartsson, H., Patz, S., 1995. The rician distribution of noisy mri data. Magnetic resonance in medicine 34, 910–914.

Hellwig, B., Schüz, A., Aertsen, A., 1994. Synapses on axon collaterals of pyramidal cells are spaced at random intervals: a golgi study in the mouse cerebral cortex. Biological cybernetics 71, 1–12.

Huang, S.Y., Tian, Q., Fan, Q., Witzel, T., Wichtmann, B., McNab, J.A., Daniel Bireley, J., Machado, N., Klawiter, E.C., Mekkaoui, C., et al., 2020. High-gradient diffusion mri reveals distinct estimates of axon diameter index within different white matter tracts in the in vivo human brain. Brain Structure and Function 225, 1277–1291.

Huang, S.Y., Tobyne, S.M., Nummenmaa, A., Witzel, T., Wald, L.L., McNab, J.A., Klawiter, E.C., 2016. Characterization of axonal disease in patients with multiple sclerosis using high-gradient-diffusion mr imaging. Radiology 280, 244–251.

Huang, S.Y., Witzel, T., Keil, B., Scholz, A., Davids, M., Dietz, P., Rummert, E., Ramb, R., Kirsch, J.E., Yendiki, A., et al., 2021. Connectome 2.0: Developing the next-generation ultra-high gradient strength human mri scanner for bridging studies of the micro-, meso-and macro-connectome. NeuroImage 243, 118530.

Jelescu, I.O., Veraart, J., Fieremans, E., Novikov, D.S., 2016. Degeneracy in model parameter estimation for multi-compartmental diffusion in neuronal tissue. NMR in Biomedicine 29, 33–47.

Jespersen, S.N., Lundell, H., Sønderby, C.K., Dyrby, T.B., 2013. Orientationally invariant metrics of apparent compartment eccentricity from double pulsed field gradient diffusion experiments. NMR in Biomedicine 26, 1647–1662.

Jespersen, S.N., Olesen, J.L., Hansen, B., Shemesh, N., 2018. Diffusion time dependence of microstructural parameters in fixed spinal cord. NeuroImage 182, 329–342.

Jian, B., Vemuri, B.C., Özarslan, E., Carney, P.R., Mareci, T.H., 2007. A novel tensor distribution model for the diffusion-weighted mr signal. NeuroImage 37, 164–176.

Johnson, V.E., Stewart, W., Smith, D.H., 2013. Axonal pathology in traumatic brain injury. Experimental neurology 246, 35–43.

Kaden, E., Kelm, N.D., Carson, R.P., Does, M.D., Alexander, D.C., 2016. Multi-compartment microscopic diffusion imaging. NeuroImage 139, 346–359.

Karlupia, N., Schalek, R.L., Wu, Y., Meirovitch, Y., Wei, D., Charney, A.W., Kopell, B.H., Lichtman, J.W., 2023. Immersion fixation and staining of multicubic millimeter volumes for electron microscopy–based connectomics of human brain biopsies. Biological Psychiatry.

Koay, C.G., Basser, P.J., 2006. Analytically exact correction scheme for signal extraction from noisy magnitude MR signals. Journal of magnetic resonance 179, 317–322.

Lampinen, B., Szczepankiewicz, F., van Westen, D., Englund, E., C Sundgren, P., Lätt, J., Ståhlberg, F., Nilsson, M., 2017. Optimal experimental design for filter exchange imaging: Apparent exchange rate measurements in the healthy brain and in intracranial tumors. Magnetic resonance in medicine 77, 1104–1114.

Lee, H.H., Fieremans, E., Novikov, D.S., 2018. What dominates the time dependence of diffusion transverse to axons: Intra-or extra-axonal water? NeuroImage 182, 500–510.

Lee, H.H., Fieremans, E., Novikov, D.S., 2021. Realistic microstructure simulator (rms): Monte carlo simulations of diffusion in threedimensional cell segmentations of microscopy images. Journal of Neuroscience Methods 350, 109018.

Lee, H.H., Jespersen, S.N., Fieremans, E., Novikov, D.S., 2020a. The impact of realistic axonal shape on axon diameter estimation using diffusion MRI. NeuroImage, 117228.

Lee, H.H., Papaioannou, A., Kim, S.L., Novikov, D.S., Fieremans, E., 2020b. A time-dependent diffusion MRI signature of axon caliber variations and beading. Communications biology 3, 1–13.

Lee, H.H., Papaioannou, A., Novikov, D.S., Fieremans, E., 2020c. In vivo observation and biophysical interpretation of time-dependent diffusion in human cortical gray matter. NeuroImage, 117054.

Lee, H.H., Yaros, K., Veraart, J., Pathan, J.L., Liang, F.X., Kim, S.G., Novikov, D.S., Fieremans, E., 2019. Along-axon diameter variation and axonal orientation dispersion revealed with 3D electron microscopy: implications for quantifying brain white matter microstructure with histology and diffusion MRI. Brain Structure and Function 224, 1469–1488.

Lovas, G., Szilágyi, N., Majtényi, K., Palkovits, M., Komoly, S., 2000. Axonal changes in chronic demyelinated cervical spinal cord plaques. Brain 123, 308–317.

Morales, J., Benavides-Piccione, R., Dar, M., Fernaud, I., Rodríguez, A., Anton-Sanchez, L., Bielza, C., Larranaga, P., DeFelipe, J., Yuste, R., 2014. Random positions of dendritic spines in human cerebral cortex. Journal of Neuroscience 34, 10078–10084.

Neuman, C., 1974. Spin echo of spins diffusing in a bounded medium. The Journal of Chemical Physics 60, 4508–4511.

Nilsson, M., Lasič, S., Drobnjak, I., Topgaard, D., Westin, C.F., 2017. Resolution limit of cylinder diameter estimation by diffusion mri: The impact of gradient waveform and orientation dispersion. NMR in Biomedicine 30, e3711.

Nilsson, M., Lätt, J., Ståhlberg, F., van Westen, D., Hagslätt, H., 2012. The importance of axonal undulation in diffusion MR measurements: a Monte Carlo simulation study. NMR in Biomedicine 25, 795–805.

Novikov, D.S., 2020. The present and the future of microstructure mri: From a paradigm shift to “normal science”. Journal of Neuroscience Methods, 108947.

Novikov, D.S., Fieremans, E., Jespersen, S.N., Kiselev, V.G., 2019. Quantifying brain microstructure with diffusion MRI: Theory and parameter estimation. NMR in Biomedicine 32, e3998.

Novikov, D.S., Jensen, J.H., Helpern, J.A., Fieremans, E., 2014. Revealing mesoscopic structural universality with diffusion. Proceedings of the National Academy of Sciences of the United States of America 111, 5088–5093.

Novikov, D.S., Kiselev, V.G., 2010. Effective medium theory of a diffusion-weighted signal. NMR in Biomedicine 23, 682–697.

Novikov, D.S., Veraart, J., Jelescu, I.O., Fieremans, E., 2018. Rotationally-invariant mapping of scalar and orientational metrics of neuronal microstructure with diffusion MRI. NeuroImage 174, 518–538.

Ronen, I., Budde, M., Ercan, E., Annese, J., Techawiboonwong, A., Webb, A.G., 2014. Microstructural organization of axons in the human corpus callosum quantified by diffusion-weighted magnetic resonance spectroscopy of n-acetylaspartate and post-mortem histology. Brain Structure & Function 219, 1773–1785.

Sepehrband, F., Alexander, D.C., Kurniawan, N.D., Reutens, D.C., Yang, Z., 2016. Towards higher sensitivity and stability of axon diameter estimation with diffusion-weighted MRI. NMR in Biomedicine 29, 293–308.

Shapson-Coe, A., Januszewski, M., Berger, D.R., Pope, A., Wu, Y., Blakely, T., Schalek, R.L., Li, P.H., Wang, S., Maitin-Shepard, J., et al., 2021. A connectomic study of a petascale fragment of human cerebral cortex. BioRxiv.

Shepherd, G.M., Raastad, M., Andersen, P., 2002. General and variable features of varicosity spacing along unmyelinated axons in the hippocampus and cerebellum. Proceedings of the National Academy of Sciences 99, 6340–6345.

Stejskal, E.O., Tanner, J.E., 1965. Spin diffusion measurements: spin echoes in the presence of a time-dependent field gradient. The journal of chemical physics 42, 288–292.

Szczepankiewicz, F., Westin, C.F., Nilsson, M., 2019. Maxwell-compensated design of asymmetric gradient waveforms for tensor-valued diffusion encoding. Magnetic resonance in medicine 82, 1424–1437.

Tandan, R., Bradley, W.G., 1985. Amyotrophic lateral sclerosis: Part clinical features, pathology, and ethical issues in management. Annals of Neurology: Official Journal of the American Neurological Association and the Child Neurology Society 18, 271–280.

Tang-Schomer, M.D., Johnson, V.E., Baas, P.W., Stewart, W., Smith, D.H., 2012. Partial interruption of axonal transport due to microtubule breakage accounts for the formation of periodic varicosities after traumatic axonal injury. Experimental neurology 233, 364–372.

Tian, Q., Ngamsombat, C., Lee, H.H., Berger, D.R., Wu, Y., Fan, Q., Bilgic, B., Novikov, D.S., Fieremans, E., Rosen, B.R., et al., 2020. Automated segmentation of human axon and myelin from electron microscopy data using deep learning for microstructural validation and simulation, in: Proc. Int. Soc. Magn. Reson. Med, p. 430.

Veraart, J., Fieremans, E., Novikov, D.S., 2019. On the scaling behavior of water diffusion in human brain white matter. NeuroImage 185, 379–387.

Veraart, J., Nunes, D., Rudrapatna, U., Fieremans, E., Jones, D.K., Novikov, D.S., Shemesh, N., 2020. Noninvasive quantification of axon radii using diffusion MRI. eLife 9, e49855.

West, K.L., Kelm, N.D., Carson, R.P., Does, M.D., 2016. A revised model for estimating g-ratio from MRI. NeuroImage 125, 1155–1158.

Westin, C.F., Knutsson, H., Pasternak, O., Szczepankiewicz, F., Özarslan, E., van Westen, D., Mattisson, C., Bogren, M., O’Donnell, L.J., Kubicki, M., et al., 2016. Q-space trajectory imaging for multidimensional diffusion mri of the human brain. Neuroimage 135, 345–362.

